# Oncogenic Notch promotes long-range regulatory interactions within hyperconnected 3D cliques

**DOI:** 10.1101/527325

**Authors:** Jelena Petrovic, Yeqiao Zhou, Maria Fasolino, Naomi Goldman, Gregory W. Schwartz, Maxwell R. Mumbach, Son C. Nguyen, Kelly S. Rome, Yogev Sela, Zachary Zapataro, Stephen C. Blacklow, Michael J. Kruhlak, Junwei Shi, Jon C. Aster, Eric F. Joyce, Shawn C. Little, Golnaz Vahedi, Warren S. Pear, Robert B. Faryabi

**Affiliations:** Department of Pathology and Laboratory Medicine, University of Pennsylvania Perelman School of Medicine, Philadelphia, PA 19104, USA; Department of Genetics, University of Pennsylvania Perelman School of Medicine, Philadelphia, PA 19104, USA; Abramson Family Cancer Research Institute, University of Pennsylvania Perelman School of Medicine, Philadelphia, PA 19104, USA; Department of Cell and Developmental Biology, University of Pennsylvania Perelman School of Medicine, Philadelphia, PA 19104, USA; Department of Genetics, Stanford University, Stanford, CA 94305, USA; Department of Biological Chemistry, Harvard Medical School, Boston, MA 02215, USA; Center for Cancer Research, National Institute of Health, Bethesda, MD 20892, USA; Department of Pathology, Brigham and Women’s Hospital, Boston, MA 02115, USA

## Abstract

Chromatin loops enable transcription factor-bound distal enhancers to interact with their target promoters to regulate transcriptional programs. Although developmental transcription factors, such as active forms of Notch, can directly stimulate transcription by activating enhancers, the effect of their oncogenic subversion on the 3-dimensional (3D) organization of the cancer genome is largely undetermined. By mapping chromatin looping genome-wide in Notch-dependent triple-negative breast cancer and B-cell lymphoma, we show that far beyond the well-characterized role of Notch as an activator of distal enhancers, Notch regulates its direct target genes through establishing new long-range regulatory interactions. Moreover, a large fraction of Notch-promoted regulatory loops forms highly interacting enhancer and promoter spatial clusters, termed “3D cliques”. Loss-and gain-of-function experiments show that Notch preferentially targets hyperconnected 3D cliques that regulate the expression of crucial proto-oncogenes. Our observations suggest that oncogenic hijacking of developmental transcription factors can dysregulate transcription through widespread effects on the spatial organization of cancer genomes.

## Introduction

Folding of chromatin into structural and regulatory chromatin loops is emerging as an important regulator of gene expression^1,2^. Chromatin-folding organization is often perturbed at different hierarchical levels in cancer^3–6^. Changes in spatial chromatin organization due to genomic rearrangements or dysregulation of conformation-associated proteins in cancer have been reported^5–10^, yet chromatin-folding reorganization in response to oncogenic subversion of developmental transcription factors, a frequent class of oncogenic drivers, is not well understood. Notch transcription complexes control cellular development and tissue homeostasis^11^ and when dysregulated contribute to the pathogenesis of multiple malignancies^12^. Here, we used Notch to examine the impact of oncogenic transcription factors on long-range chromatin contacts among and between enhancers and promoters in Notch-dependent tumors.

Notch target genes play crucial oncogenic roles in several hematologic malignancies and solid tumors^12^. Activating Notch mutations often disrupt the Notch negative regulatory region (NRR) or C-terminal PEST degron domain, producing ligand-independent release of the Notch intracellular domain (NICD) or an increase in NICD half-life, respectively. NICDs translocate to the nucleus and form Notch transcription complexes (NTCs) with the DNA-binding factor RBPJ and other co-factors. Oncogenic Notch transcription complexes recruit histone acetyltransferase p300^13^, histone demethylase KDM1A^14,15^ and components of the mediator complex^16^ to Notch-responsive elements to turn on the transcription of target genes with oncogenic activity. In hematologic malignancies, Notch binding events are often associated with increased histone acetylation and activation of distal enhancer elements^17^. Direct regulation of the proto-oncogene *MYC* in both B- and T-lymphoid malignancies by Notch-activated enhancers, which are located up to 1.5 Mb away from the *MYC* promoter, exemplifies Notch-dependent long-range gene regulation^18–21^. Although looping of chromatin, which enables physical contacts between Notch-bound enhancers and promoters, is essential for proper and selective gene expression, it remains unclear to what extent Notch transcription complexes influence long-range regulatory contacts.

Chromatin loops, juxtaposing transcription factor-bound distal enhancers with the promoters of target genes are facilitated by structural proteins, including the DNA-binding insulator protein CCCTC-binding factor (CTCF) and cohesin^5–8,22–25^. Ring-shaped cohesin complexes are loaded at active enhancers and promoters to stabilize their physical interactions^26–29^. Enhancer-promoter loops are mostly constrained within larger genome organizational structures, variably referred to as contact domains, interaction domains, topologically associated domains (TADs), sub-TADs, loop domains, and insulated neighborhoods^1,4,28,30–34^, the boundaries of which are occupied by cohesin complexes and CTCF^1,6,28,30,32–35^. More recently, it was shown that the ubiquitous transcription factor YY1, in addition to a limited number of architectural proteins, binds to enhancers and facilitates their looping to promoters, suggesting that enhancer-promoter interactions could be mediated by particular transcription factors bound at DNA elements engaged in transcriptional regulation^36^.

Oncogenic Notch transcription complexes bind distal enhancers^17,18^, raising the question of whether oncogenic Notch regulates transcription by positively influencing interactions among and between enhancers and promoters. To investigate the impact of oncogenic Notch on the 3D genome organization of cancer cells, we generated cohesin HiChIP and 1D epigenomic data sets in two different Notch-dependent cancer cell types, triple-negative breast cancer (TNBC) and mantle cell lymphoma (MCL), in the Notch-on and -off states. We report here that Notch transcription complexes control their direct target genes through two distinct regulatory modes: either through existing loops or by facilitating new long-range regulatory interactions. This combination of pre-existing and Notch-promoted loops coalesce enhancers and promoters to form highly interacting clusters, termed “3D cliques”. Notch preferentially activates enhancers and promotes looping interactions within highly connected 3D cliques that regulate key oncogenes. These observations suggest a general mechanism that oncogenic transcription factors can exploit to regulate the transcriptional outputs of cancer cells.

## Results

### Contact domains of Notch-mutated cancer cells are lineage invariant

Genome-wide chromatin conformation capture methods such as HiChIP, PLAC-seq and ChIA-PET were used to accurately map chromatin contact domains in multiple cell types^30,37–40^. Given that the cohesin complex is loaded at enhancer-promoter loops and is involved in CTCF-mediated interactions, we first performed HiChIP for the cohesin subunit SMC1a in MCL Rec-1 cells with an activating Notch mutation and used chromosome-wide insulation potential^41^ to systematically identify high-confidence contact domain boundaries (Table S1). To validate the sensitivity and specificity of our cohesion HiChIP, we compared the Rec-1 contact domains with the contact domains delineated by *in situ* Hi-C in the EBV-transformed GM12878 lymphoblastoid B cell line^33^. GM12878 cells express an EBV-encoded RBPJ-binding factor, EBNA2, that mimics Notch activities^42^, and its genome organization is similar to Rec-1 cells^18^. The contact domain boundaries identified in Rec-1 cells using cohesin HiChIP (~760 million sequenced reads) were concordant with the ones identified by *in situ* Hi-C of GM12878 cells^33^ (~3 billion sequenced reads), both at the level of a single chromosome (Figure S1A) and genome-wide (Fisher’s exact p-value < 1E-15, Figure S1B). This level of reproducibility was similar to that observed when the HiChIP^37^ and *in situ* Hi-C of GM12878 were compared (Fisher’s exact p-value < 1E-04, Figure S1C). These results demonstrate that our cohesin HiChIP data were of high quality and provide an efficient method to accurately delineate chromatin contact domains with ~4-fold lower sequencing depth in Notch-mutated cells.

In addition to MCL, activating Notch mutations are frequent in TNBC^43–47^. We performed cohesin HiChIP in the Notch-mutated TNBC cell lines HCC1599 and MB157^44,46^, analyzing more than 1.5 billion read pairs. A sizable fraction of contact domains is enclosed by a structural chromatin loop with CTCF-bound and cohesin-occupied anchors^1,28,48–50^. Examination of CTCF and cohesin binding events showed that these proteins co-occupy 81.4% of MB157 contact domain boundaries (proportion test p-value < 1E-15, Figure S1D). As expected for a high-quality data set, more than 85% of the CTCF-bound contact domain boundaries in MB157 cells had inward-oriented CTCF motifs (785 inward-oriented versus 2 outward-oriented, Figure S1E).

TNBC HCC1599 and MB157 contact domains showed highly similar organization, as exemplified by the organization of chromosome 8 (Figure 1A and Table S1). Genome-wide, out of 4,767 and 4,847 contact domain boundaries identified in HCC1599 and MB157 cells, respectively, 4,223 domain boundaries were common to both (Fisher’s exact p-value < 1E-15, Figure S1F). Furthermore, most contact domain boundaries were also shared by MCL Rec-1 and TNBC cells, as exemplified here by *MYC* locus at chromosome 8 (Figure 1B, 5 Kb genomic resolution). Genome-wide, nearly 70% of the identified contact domain boundaries were shared by these two different Notch-mutated cancer cell lineages (Fisher’s exact p-value < 1E-15, Figure 1C), consistent with the extent of concordance previously noted when other cell lineages were compared^1,32,33,51^. Thus, chromatin contact domains are largely lineage-independent organizational features of Notch-mutated cancer cell genomes.

**Figure 1:**
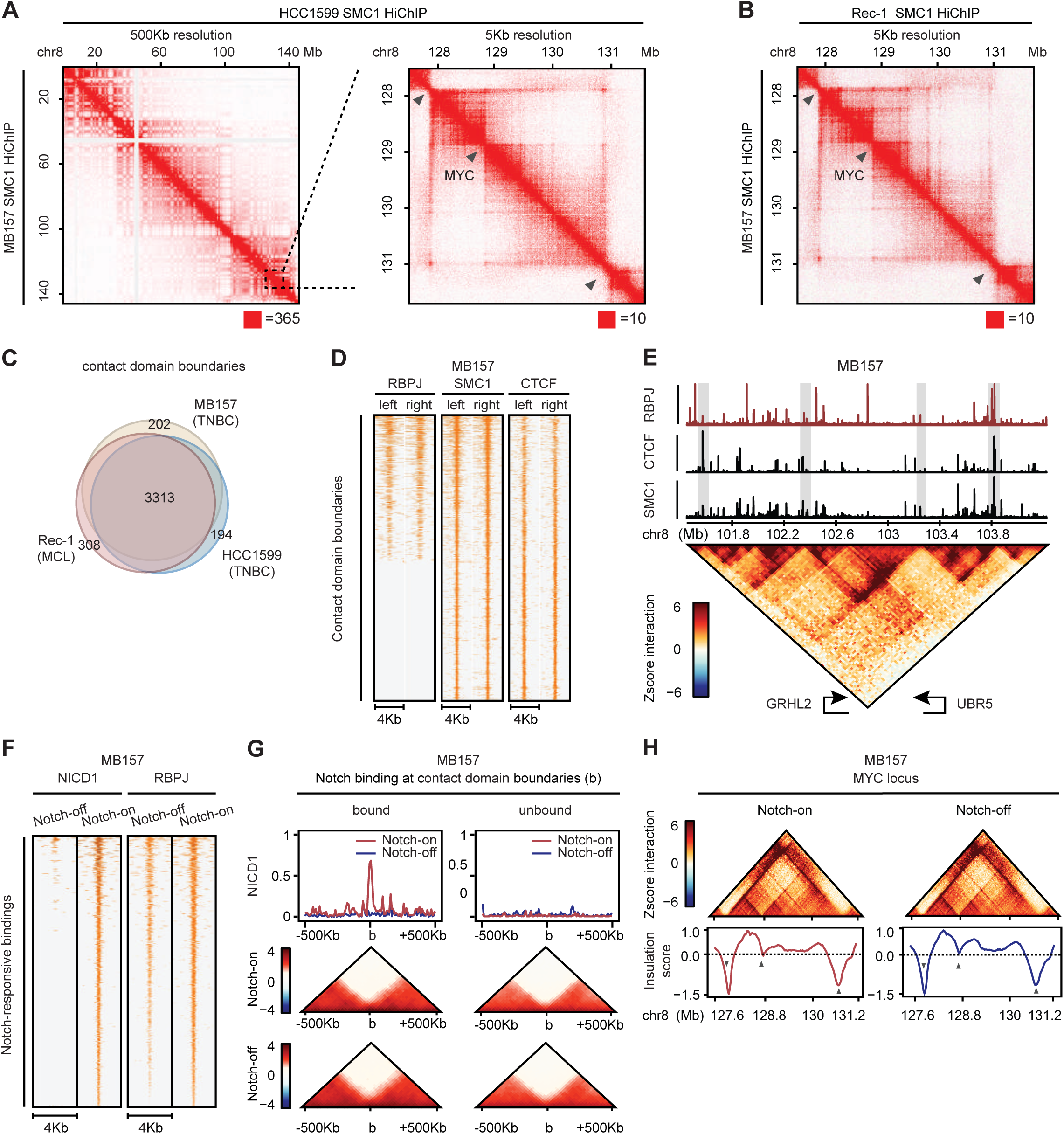
Contact domains of Notch-mutated tumors are lineage and Notch signaling insensitive. (A) Contact matrices showing MB157 SMC1 HiChIP, lower half, and HCC1599 SMC1 HiChIP, upper half, share contact domain boundaries at chromosome 8. Left: the whole chromosome, at 500 Kb resolution; right: 127.5-131.5 Mb *MYC* locus shown at 5 Kb resolution. Red square on the bottom right of each panel indicates maximum intensity. Gray arrows: boundaries demarcated by local minimum detection of insulation score. (B) Contact matrices showing MB157 SMC1 HiChIP, lower half, and Rec-1 SMC1 HiChIP, upper half, share contact domain boundaries at chr8: 127.5-131.5 Mb *MYC* locus shown at 5 Kb resolution. Gray arrows: boundaries demarcated by local minimum detection of insulation score. (C) Venn diagram comparing the contact domain boundaries of Rec-1, HCC1599 and MB157 depicts that a significant number of boundaries are shared (Fisher’s exact p-value < 1E-15). (D) Heatmap displaying RBPJ, SMC1 and CTCF occupancy on left and right boundaries of 920 MB157 contact domains. (E) Contact map (bottom) and genome-browser tracks of RBPJ, CTCF and SMC1 ChIP-seq at *GRHL2* locus showing contact domain boundaries are enriched for RBPJ, CTCF and SMC1 binding. The intensity of each pixel on the 25 Kb-binned contact map represents the Z-score transformed interaction frequency between two loci. Gray box: RBPJ, CTCF and SMC1 binding events on contact domain boundaries. (F) Heatmap of NICD1 and RBPJ occupancy showing 3,216 reproducible Notch-responsive elements determined with IDR pipeline in MB157 with a significant decrease (enrichR FDR < 0.05) in Notch-off (GSI) versus Notch-on (GSI-washout). (G) Metagene analyses (top) showing Notch occupancy, and pile-up plots (bottom) depicting aggregated Z-score interaction on MB157 domain boundaries in Notch-on (DMSO) and Notch-off (GSI) conditions where the overall differential boundary insulation scores are insignificant (Wilcoxon rank sum test p-value > 0.15). Left: centered around 1,003 Notch-bound domain boundaries. Right: matching number of Notch-unbound boundaries. (H)Contact map (top) and insulation profile (bottom) at *MYC* locus showing contact domain boundaries are unaltered in Notch-on (DMSO) and Notch-off (GSI) conditions. Gray arrows: boundaries demarcated by local minimum detection of insulation score.

### Contact domains are insensitive to Notch signals

To test the effect of Notch transcription complexes binding on genome organization, we first performed RBPJ ChIP sequencing (ChIP-seq) in MB157. We observed 19% of the RBPJ binding events localized to domain boundaries (Figure S1G) and, conversely, 43% of CTCF-bound, cohesion-occupied boundaries showed significant RBPJ binding (permutation proportion 14%, proportion test p-value < 1E-15, Figures 1D and 1E). To determine if Notch transcription complex binding influenced insulation potential of contact domain boundaries, we performed a gamma-secretase inhibitor (GSI)-washout assay^52^, which permits timed loading of Notch transcription complexes onto chromatin^17^. Notch transcription complex binding to genomic elements involved in Notch target gene regulation exhibit rapid loading following GSI washout and Notch activation^17^, an event that also increases the RBPJ occupancy at regulatory sites^17,53^. By comparing the Notch active (GSI-washout, Notch-on) and inactive (GSI, Notch-off) states in MB157 cells, we identified 3,216 Notch-response elements with significant and reproducible increase in NICD1 and RBPJ occupancy (Figure 1F, Table S2). However, Notch-and RBPJ-bound contact domain boundaries were, on average, unaffected by Notch activity, as shown by pile-up plots of interactions centered on contact domain boundaries (Figure 1G). This observation was confirmed by inspection of genomic-distance-adjusted chromatin interaction maps and insulation profiles of the *MYC* locus before and after Notch inhibition (Figure 1H). Measurements of Notch transcription complex binding events in HCC1599 TNBC cells also showed that while Notch transcription complexes bound to many HCC1599 contact domain boundaries (Figures S1H and S1I), alterations in Notch activity did not impact contact domain integrity (Figures S1J and S1K). Together, these data suggest that contact domains are unaffected by the presence or absence of Notch transcription complex binding.

### Chromatin state of contact domains in Notch-mutated tumors is lineage-specific

Contact domains generally restrict propagation of chromatin states along chromosomes^1,30,54^. In line with this prediction, contact domains in MB157 and HCC1599 cell lines were either enriched for active (H3K27ac and/or H3K4me1 histone marked) or repressed (H3K27me3 marked) chromatin (Figures 2A, 2B, and S2A-D), with active contact domains containing, on average, twice the number of expressed genes as repressed domains. Repressed contact domains with higher H3K27me3 were on average 1.37 Mb and larger than active contact domains whose average size was 0.92 Mb (Figures 2B and S2C). Overall, the chromatin states in 79% of contact domains were identical in MB157 and HCC1599 TNBC cell lines (Fisher’s exact p-value < 1E-15, Figure S2E). Analysis of active and repressed contact domains in Notch-mutated MCL cells (Figures S2F-H) and comparison with TNBC cells showed that on average 37% of contact domain chromatin states were lineage-specific (Fisher’s exact p-value < 1E-06, Figure 2C). Together, our data indicate that while contact domains are largely invariant in Notch-mutated MCL and TNBC, the chromatin signature within contact domains is lineage-specific.

**Figure 2.**
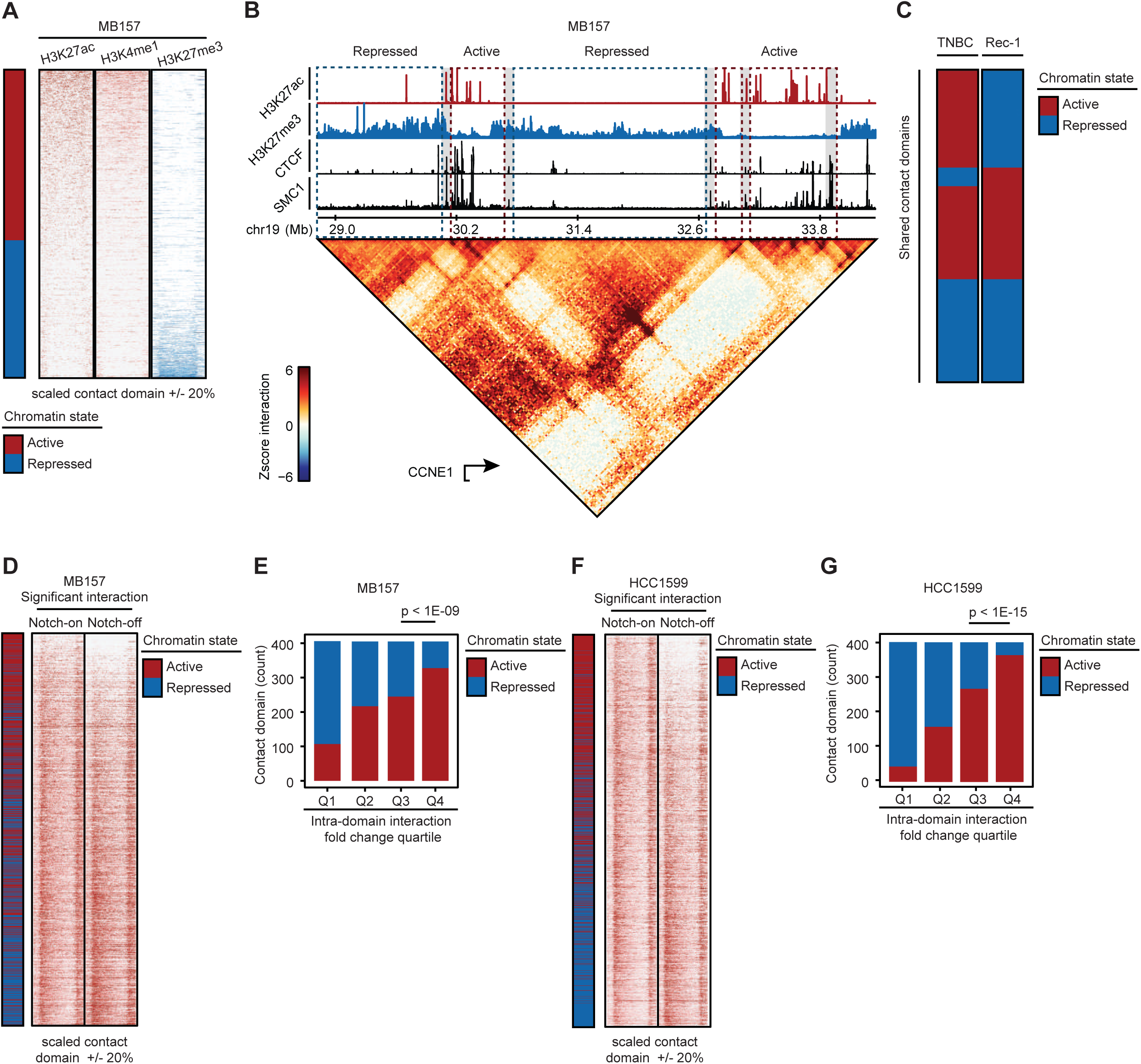
Notch sensitivity of chromatin contacts within TNBC active enhancer-marked contact domains. (A) Heatmap displaying normalized H3K27ac, H3K4me1 and H3K27me3 levels within 1,621 contact domains with significant intradomain contacts in MB157. Each domain is categorized into active or repressed based on the differential H3K27ac and H3K27me3 total level and sorted in descending order. (B) Contact map (bottom) and genome browser tracks (top) in MB157 showing that H3K27ac and H3K27me3 marks are insulated within contact domain boundaries demarcated by CTCF and SMC1 at *CCNE1* locus. Red/blue dashed-line boxes: active/repressed domains. Gray boxes: CTCF and SMC1 binding events on the boundaries. (C) Heatmap showing chromatin state of contact domains shared between TNBC (HCC1599 and MB157) and MCL (Rec-1) cell lines. (D) Heatmap of normalized significant interactions at scaled and flanked contact domains in MB157 in Notch-on (DMSO) and Notch-off (GSI) conditions. Contact domains are ranked by change (log2 fold change) of total intradomain contacts with the chromatin states indicated on the left. The overall differential intradomain contact frequency is significant (paired t-test p-value < 1E-15). (E) Barplot depicting the number of active/repressed contact domains per quartile of total intradomain contact frequency change in MB157 (proportion test p-value < 1E-09). (F) Heatmap as (D) in HCC1599 (paired t-test p-value < 1E-15). (G) Barplot as (E) in HCC1599 (proportion test p-value < 1E-15).

### Interactions within active enhancer-marked contact domains are Notch sensitive in TNBC

Intradomain interactions linking regulatory DNA elements, such as enhancers and promoters, are implicated in gene control^26,30,31,55^. To determine whether Notch signaling impacts intradomain interactions in TNBC, we first used 286 million unique read pairs of MB157 cohesin HiChIP to identify high-resolution (~5 Kb) significant interactions. We relied on a statistical model that controls for both the protein occupancy level and linear genomic distance between the connected DNA loop anchors^56^. This approach identified 265,216 significant cohesin-associated DNA interactions in MB157 cells supported by at least 4 read pairs (Table S3). Unless stated otherwise, the high-confidence set of interacting loci (also referred to as significant interactions) was used for further quantitative analysis. After Notch inhibition in MB157 cells, 236 contact domains showed at least a 4-fold decrease in overall intradomain interaction (Figure 2D). These Notch-sensitive contact domains were enriched within the active chromatin state (proportion test p-value < 1E-09, Figure 2E). To independently confirm this observation, we studied HCC1599 cells where 472,073 significant cohesin-associated DNA interactions were identified (Table S3). We again detected Notch-sensitivity of intradomain interactions connecting loci within contact domains with high loads of active enhancer histone marks (proportion test p-value < 1E-15, Figures 2F and 2G). Together, these results show that in contrast to invariant contact domain boundaries, long-range intradomain chromatin loops with potential regulatory functions are Notch-sensitive in TNBC.

### Notch activates TNBC distal enhancers

We next quantitated the direct effect of Notch on active enhancers in TNBC MB157 and HCC1599 cells. On average, Notch binding events were 19 Kb away from the closest transcription start site (Figure S3A). H3K27ac, a histone mark of active enhancers, was deposited at nearly 85% of the Notch transcription complex-bound chromatin (proportion test p-value < 1E-15, Figure S3B). Furthermore, Notch inhibition markedly decreased the H3K27ac levels, while having negligible effects on the level of H3K4me1 (Figure S3B). Together, these data suggest that Notch transcription complexes preferentially bind and activate distal enhancers in TNBC, as in T-ALL and MCL^17,18,20^

### Notch-promoted and preformed enhancer-promoter contacts regulate direct Notch target genes in TNBC

Regulatory DNA loops between promoters and distal enhancers are crucial for proper gene control^57,58^. We thus asked whether distal Notch-responsive elements (Figures S3A and S3B) that are likely to directly regulate TNBC transcriptional outputs are also associated with Notch-sensitive intradomain long-range interactions (Figures 2E and 2G). To this end, we first identified TNBC Notch-sensitive genes (i.e. Notch-upregulated genes) using RNA-seq in MB157 and HCC1599 cells. Notch activation concordantly increased the transcription level of 2,038 genes in these two cell lines (Table S4). To assess the lineage-specificity of Notch-sensitive genes, we also performed differential gene expression analysis in T-ALL DND41 and MCL Rec-1 cells. We found that 504 genes, including the well-characterized Notch target genes *MYC*, *HES1*, and *CR2*^15,18,44^, were positively regulated by Notch in all three Notch-mutated cell types (Figure S3C, Table S4). Lineage-independent Notch-activated genes were enriched for known MYC targets and MYC-regulated biological processes (Figure S3D), suggesting that these genes predominantly belong to a MYC-driven expression program and are a secondary effect of Notch activation in these three Notch-mutated cancer cell types. Overall, 204 Notch sensitive genes were specific to TNBC (Figure S3C, Table S4), including *CCND1*, *KIT* and *SAT1*, all of which have been implicated in TNBC^43,59–62^. Furthermore, gene set enrichment analysis showed the TNBC-specific Notch target genes were enriched for genes associated with breast cancer and mammary epithelium biology (Figures S3C and S3E, Table S5). These data indicate the existence of a TNBC-specific Notch-driven transcription program, in line with the lineage-specificity of the TNBC contact domain chromatin state (Figure 2C).

We next assessed how Notch-dependent gene expression relates to TNBC 3D genome organization. After aligning the Notch-sensitive genes to the TNBC 3D genome landscape, we observed that these genes were preferentially associated with active contact domains and with Notch-sensitive intradomain interactions (Fisher’s exact p-value < 1E-04, Figure 3A). Integrative analysis of Notch-binding events, Notch-dependent enhancers (Table S6) and transcripts, and high confidence chromatin loops distinguished direct from indirect Notch targets, and identified 215 and 386 direct Notch-upregulated genes (i.e. Notch-activated genes) in MB157 and HCC1599 cells, respectively. In both TNBC lines, inhibition of Notch signaling markedly decreased H3K27ac level at Notch-bound enhancer elements linked to Notch-activated genes (Wilcoxon rank sum p-value < 1E-15, Figure 3B). We also observed a significant reduction in the frequency of long-range interactions between the Notch-bound DNA elements and their target promoters upon Notch inhibition (Wilcoxon rank sum p-value < 1E-03, Figure 3C).

**Figure 3.**
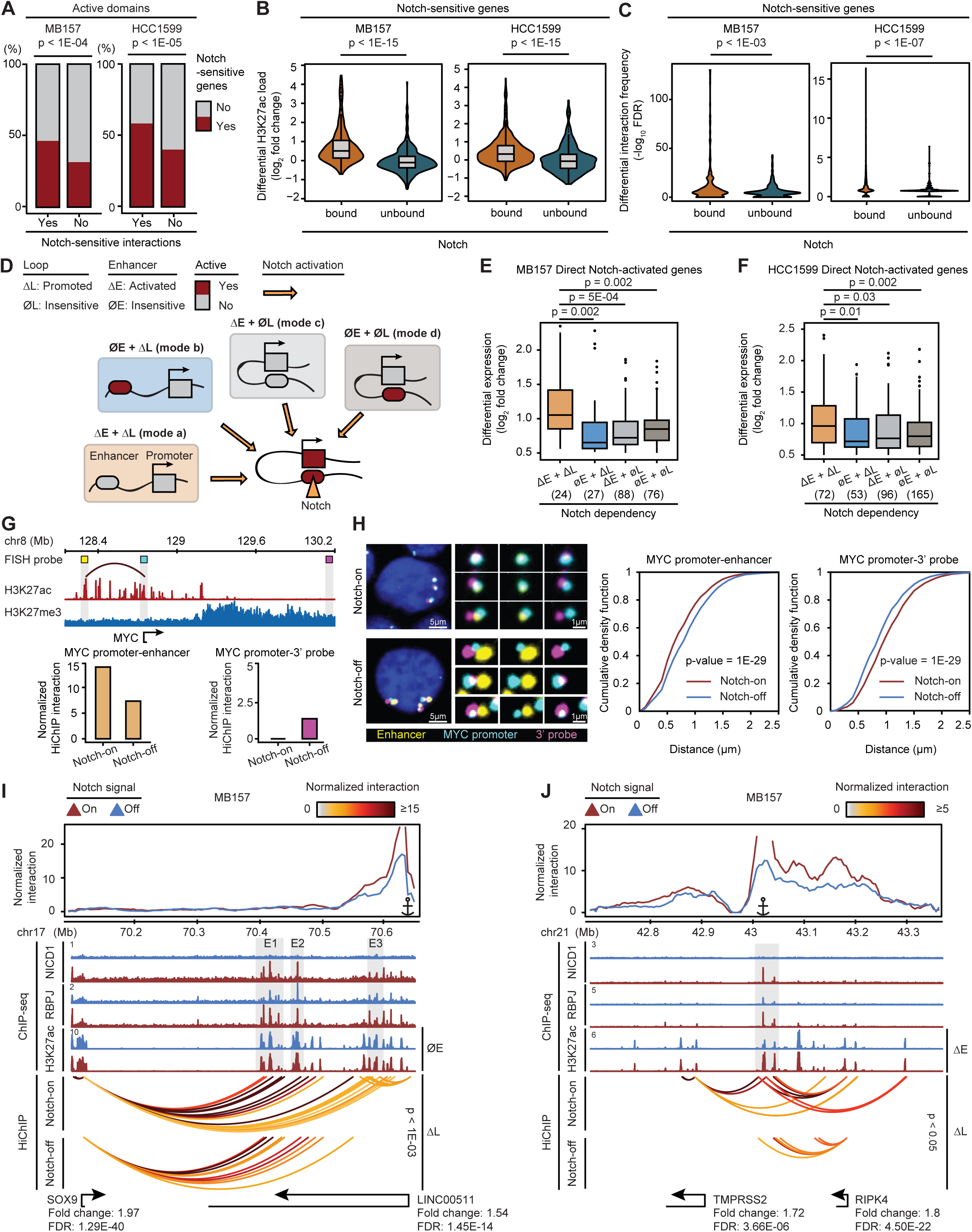
Activation of direct Notch target genes by Notch-promoted and preformed enhancer-promoter contacts in TNBC. (A) Barplots showing the proportion of contact domains encompassing any Notch-sensitive gene conditioned on whether intradomain interactions were Notch-sensitive or Notch-insensitive in MB157 (Fisher’s exact test p-value < 1E-04) (left) and HCC1599 (Fisher’s exact p-value < 1E-05) (right). (B) Box and violin plots of differential (log2 fold change) H3K27ac level at Notch-bound or -unbound distal enhancers of Notch-sensitive genes in MB157 (left) and HCC1599 (right) (Wilcoxon rank sum p-value < 1E-15). (C) Violin plots of differential (-log_10_ FDR) frequency of interactions between Notch-bound or unbound distal enhancers and Notch-sensitive gene promoters in MB157 (left, Wilcoxon rank sum p-value < 1E-03) and HCC1599 (right, Wilcoxon rank sum p-value < 1E-07). (D) Model depicting four possible Notch regulatory modes controlling Notch direct target genes by combinations of Notch-bound and -promoted loops (ΔL), Notch-bound and -activated enhancers (ΔE), Notch-bound but Notch-insensitive loops (ØL), and Notch-bound but Notch-insensitive enhancers (ØE). (E) Boxplot showing differential (log_2_ fold change) gene expression of direct Notch-activated genes in MB157 categorized by Notch-dependency of interacting Notch-bound enhancers and loops per mode in (D). Notch-bound and -promoted loops (ΔL), Notch-bound and -activated enhancers (ΔE), Notch-bound but Notch-insensitive loops (ØL), and Notch-bound but Notch-insensitive enhancers (ØE). Number of genes in each mode is listed in parenthesis. (F) Boxplot as in (E) for HCC1599 line. (G) Frequency of interactions between the *MYC* promoter and enhancer significantly decreased after Notch inhibition. Top: ChIP-seq tracks showing H3K27ac and H3K27me3 load at *MYC* locus marked with matching colors for position of the probes against the *MYC* promoter (cyan), *MYC* enhancer (yellow), and H3K27me3-marked *MYC* 3’ (magenta) sequences for a 3-color DNA FISH. Bottom: The quantification of HiChIP-measured contact frequency between the *MYC* promoter and *MYC* enhancer probes (left), and *MYC* promoter and *MYC* 3’ probes (right) in Notch-on (DMSO) and Notch-off (GSI) conditions in MB157 cells are compared. (H) Distances between the *MYC* promoter and enhancer significantly increased after Notch inhibition in MB157 cells. Left: examples of cells and magnified images for 3-color, cyan-yellow, and cyan-magenta, from left to right, in Notch-on (DMSO) and Notch-off (GSI) in MB157 cells are shown. Blue: DAPI. Scale bars: Left: 5 µm; Right: 1 µm. Center: Cumulative density functions of distances between *MYC* promoter and the closest *MYC* enhancer probe in the same cells are compared between Notch-on cells (red, N = 4,314) and Notch-off cells (blue, N = 5,271). Mean (+/-S.D.) distance of Notch-on and Notch-off are 0.71 +/-0.30 µm, and 0.83 +/-0.32 µm, respectively (Kolmogorov-Smirnov test p-value = 1E-29). Left: Cumulative density functions of distances between *MYC* promoter and the closest *MYC* 3’ probe in the same cells are compared between Notch-on and Notch-off cells. Mean (+/-S.D.) distance of Notch-on and Notch-off were 1.0 +/-0.32 µm, and 0.88 +/-0.31 µm, respectively (Kolmogorov-Smirnov p-value = 1E-29). (I) Notch-promoted loops (ΔL) linking Notch-insensitive active enhancers (ØE) E1 and E2 to *SOX9* promoter and E3 to *LINC00511* promoter in MB157. Top panel: virtual 4C plot depicting the normalized interaction frequency from *LINC00511* promoter viewpoint. ChIP-seq tracks showing Notch-sensitive NICD1 and RBPJ occupancy, and Notch-insensitive H3K27ac load marked with gray boxes. HiChIP arcs displaying normalized significant interactions of *SOX9* and *LINC00511* promoters to distal enhancers in Notch-on (top, DMSO) and Notch-off (bottom, GSI) (paired t-test p-value < 1E-03). Bottom track indicating *SOX9* and *LINC00511* Ensembl gene positions and their expression fold change and FDR determined by DESeq2. (J) Notch-promoted loops (ΔL) enabling spatial co-regulation of *TMPRSS2* and *RIPK4* genes in TNBC through shared Notch-activated enhancers (ΔE). ChIP-seq tracks showing Notch-sensitive NICD1 and RBPJ occupancy and H3K27ac load marked with gray box. HiChIP arcs displaying normalized significant interactions between enhancers and promoters in Notch-on (top, DMSO) and Notch-off (bottom, GSI) in MB157 (paired t-test p-value < 0.05). Bottom track indicating Ensembl gene positions and their expression fold change and FDR determined by DESeq2.

We defined Notch-promoted loops as Notch-sensitive interactions that connect promoters to distal Notch transcription complex-bound elements. To further dissect the mechanisms of direct gene activation by Notch transcription complexes, we considered four possible regulatory modes: (a) Notch-promoted loops (ΔL) linking Notch-activated enhancers (ΔE) to promoters; (b) Notch-promoted loops (ΔL) linking Notch-independent active enhancers (ØE) to promoters; (c) Notch-independent (preformed) loops (ØL) linking Notch-activated enhancers (ΔE) to promoters; (d) Notch-independent (preformed) loops (ØL) linking Notch-independent active enhancers (ØE) to promoters (Figure 3D). Integrating the Notch-dependent regulatory loops, enhancers, and Notch-binding events from MB157 cells showed that the greatest increase in transcription between the Notch-off and -on states occurred in genes in which Notch both activated the enhancers and promoted enhancer-promoter interactions (mode a, Wilcoxon rank sum p-value < 2E-03, Figure 3E). Similar observations were made in HCC1599 cells (mode a, Wilcoxon rank sum p-value < 3E-02, Figure 3F). Although “mode a” was associated with a more pronounced effect on expression of direct Notch gene targets, our analysis also identified a group of Notch-activated genes in which only loops (mode b) or enhancers (mode c) were Notch-dependent (Figure 3D). Finally, transcriptional outputs of another group of genes linked to Notch-independent enhancers through preformed loops were shown to only depend on distal Notch binding, suggesting that Notch functions as the final transcriptional trigger at these loci (mode d, Figures 3D-F and S3F).

We next closely scrutinized Notch-promoted enhancer-promoter contacts (Figure 3D modes a and b), a previously unrecognized mode of Notch-dependent gene regulation. The proto-oncogene *MYC*, a known Notch direct target in B-and T-lymphoid malignancies^18–21^, exemplifies genes with Notch-promoted enhancer-promoter loops in TNBC (Figure 3G, also see Figure 5). To independently evaluate the Notch-dependency of promoter-enhancer loop formation at this critical proto-oncogene, we performed 3D DNA-fluorescence in situ hybridization (FISH) for three loci in MB157 cells: 1) the *MYC* promoter; 2) a *MYC* enhancer located 451 Kb 5’ of the promoter that interacted with the promoter through a Notch-sensitive long-range interaction; 3) an H3K27me3-marked T-ALL-specific Notch-dependent *MYC* enhancer located 3’ of the *MYC* promoter which was inactive in TNBC. In concordance with the HiChIP-measured decrease in *MYC* promoter-enhancer interaction frequency (Figure 3G), FISH analysis showed that the *MYC* promoter and the *MYC* 5’ enhancer probes became significantly separated upon Notch inhibition (Figure 3H). Interestingly, we observed that the *MYC* promoter and the 3’ *MYC* probes became markedly closer after Notch inhibition (Figure 3H), as observed throughout TNBC genomes (Figures 2E and 2G). Critically, for both cases the FISH data agreed with changes seen in the HiChIP-measured contact frequencies (Figures 3G and 3H). Together, these data support the observation of Notch-promoted long-range interactions in TNBC as measured by cohesin HiChIP.

**Figure 4.**
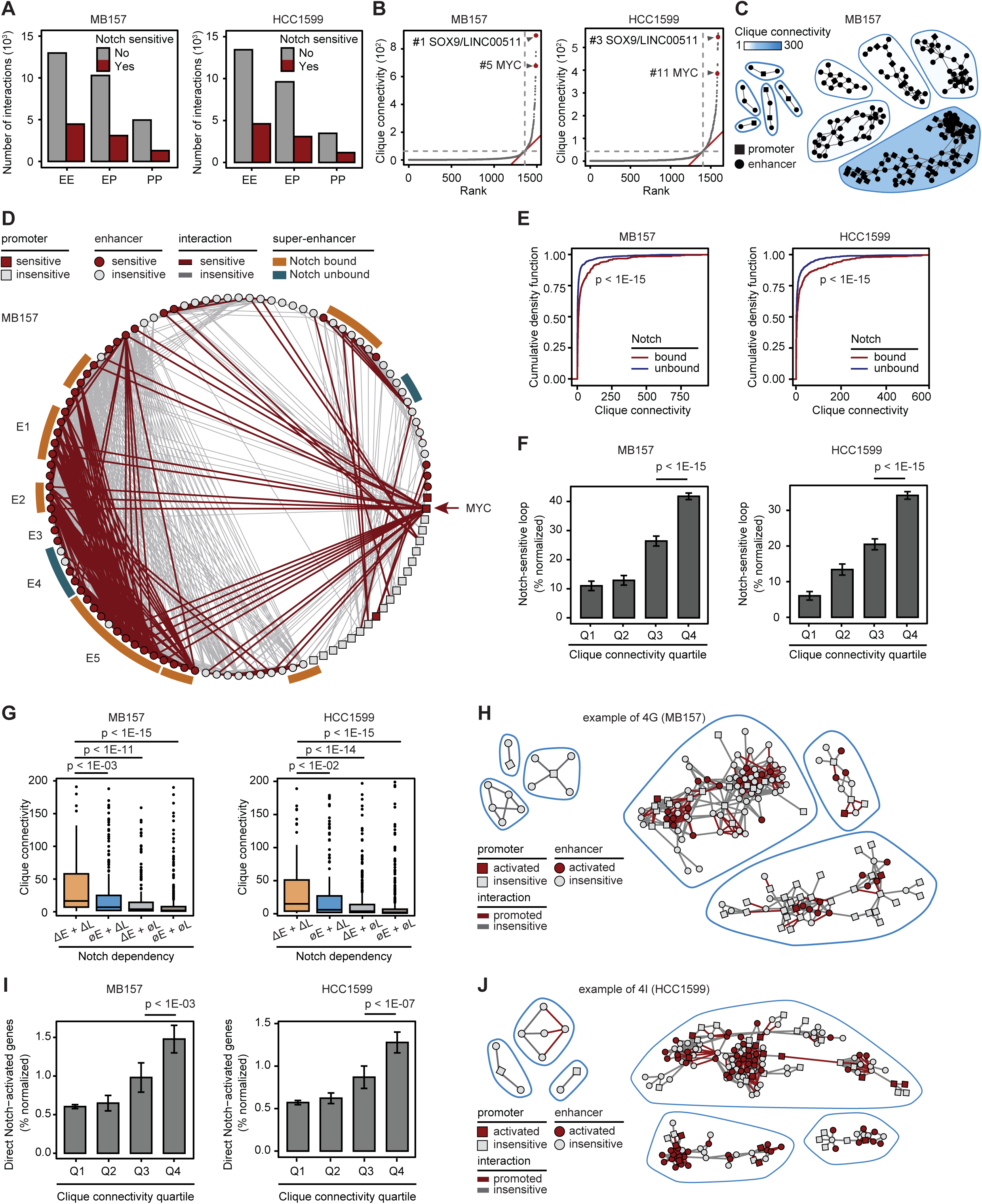
Notch targets TNBC’s hyperconnected 3D regulatory cliques. (A) Barplots showing the number of Notch-sensitive and -insensitive enhancer-enhancer (EE), enhancer-promoter (EP), promoter-promoter (PP) interactions in MB157 (left) and HCC1599 (right). (B) Distribution of 3D cliques connectivity revealing two classes of interacting enhancers and promoters. Cliques are plotted in an ascending order of their total connectivity for MB157 (left) and HCC1599 (right). Hyperconnected cliques are defined as the ones above the elbow of 3D clique total connectivity distribution. Example 3D cliques are marked and named with their representative Notch-sensitive genes. (C) Five randomly selected cliques from either below (left) or above (right) the elbow point of the curve in (B) demonstrating the asymmetric distribution of clique total connectivity in MB157. (D) Circos plot showing the clique associated with *MYC* in MB157. Red-marked circle (square) and line depicting Notch-sensitive enhancer (promoter) and significant long-range interactions, respectively. E1 to E5 mark groups of enhancers in descending linear genomic distance to *MYC* promoter located within the *MYC* 5’ contact domain. (E) Cumulative density function of clique total connectivity with or without Notch-bound enhancers in MB157 (left) and HCC1599 (right) (Wilcoxon rank sum p-value < 1E-15). (F) Barplots depicting the average ± SEM corrected percentage of Notch-sensitive loops per quartile of clique total connectivity distribution in MB157 (left) and HCC1599 (right) (Wilcoxon rank sum p-value < 1E-15). (G)Boxplots showing the total connectivity of cliques containing promoters associated with combination of Notch-bound and -promoted loops (ΔL), Notch-bound and -activated enhancers (ΔE), Notch-bound but Notch-insensitive loops (ØL), and Notch-bound but Notch-insensitive enhancers (ØE) in MB157 (left) and HCC1599 (right). (H)Three randomly selected cliques in MB157 associated with either Notch-insensitive enhancers and loops (ØE+ØL, left) or Notch-activated enhancers and Notch-promoted loops (ΔE+ΔL, right) emphasizing on the difference in clique total connectivity. (I)Barplots depicting the average ± SEM corrected percentage of direct Notch-activated genes per quartile of clique total connectivity distribution in MB157 (left, Wilcoxon rank sum p-value < 1E-03) and HCC1599 (right, Wilcoxon rank sum p-value < 1E-07). (J)Three randomly selected cliques in either the first (left) or fourth (right) quartile of clique total connectivity distribution highlighting the difference in the number of direct Notch-activated genes.

**Figure 5.**
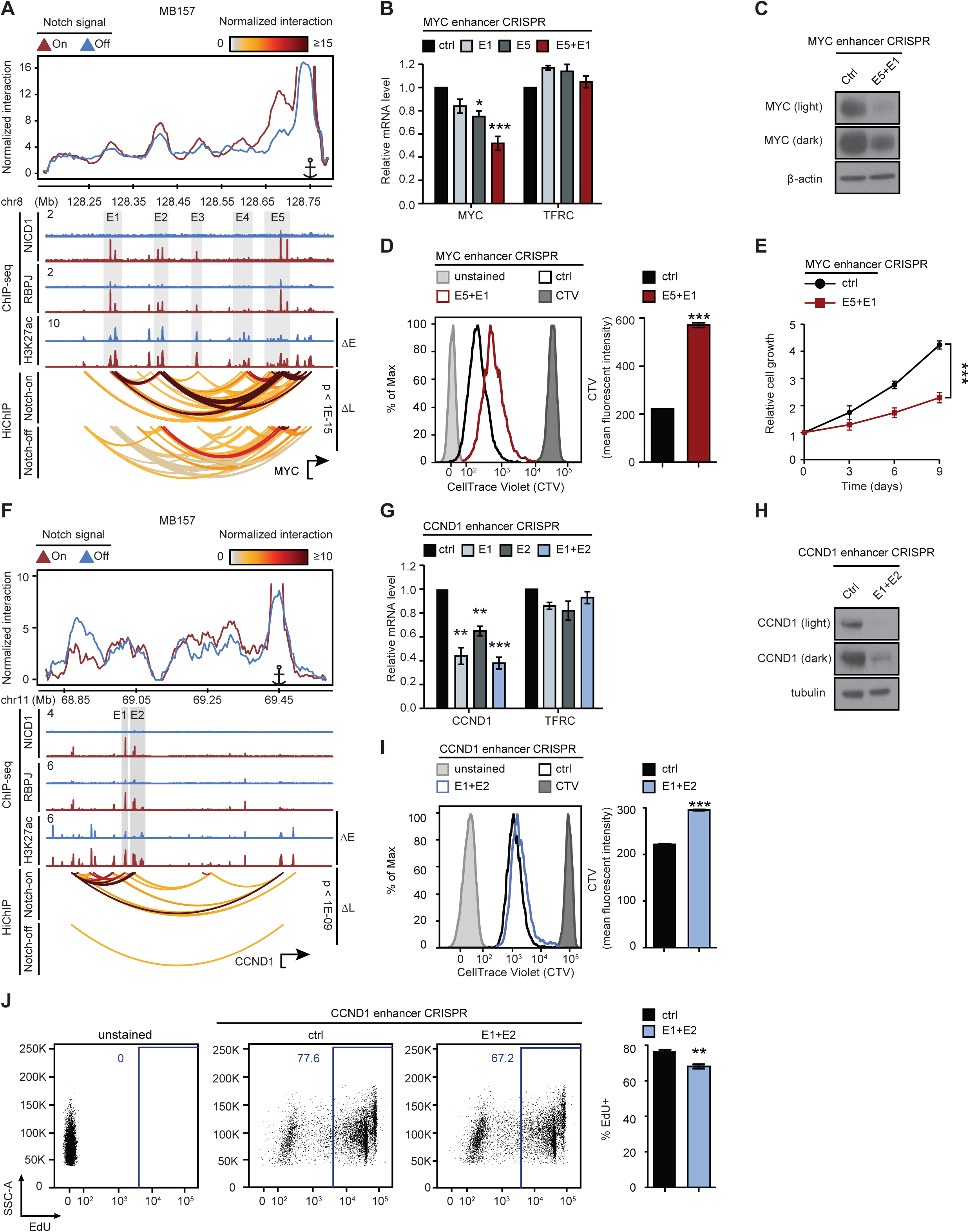
*MYC* but not *CCND1* Notch-activated distal enhancers cooperate to increase gene expression. (A) Notch-promoted loops (ΔL) linking Notch-activated enhancers (ΔE) to *MYC* in TNBC MB157 line. Top panel: virtual 4C plot depicting the normalized interaction frequency from *MYC* promoter viewpoint. ChIP-seq tracks showing Notch-sensitive NICD1 and RBPJ occupancy, and Notch-sensitive H3K27ac level marked with gray box. HiChIP arcs displaying normalized significant interactions between *MYC* promoter and distal enhancers in Notch-on (top, DMSO) and Notch-off (bottom, GSI) (paired t-test p-value < 1E-15). Bottom track indicating *MYC* Ensembl gene position. (B) Barplots showing qRT-PCR measurements of *MYC* mRNA after transduction of Cas9-expressing MB157 cells with control sgRNAs, sgRNAs targeting the E1, E5 or both reveal cooperativity of Notch-sensitive E1 and E5 *MYC* enhancers. TFRC: negative control. Data represent mean ± SEM of n=3-7 independent experiments. t-test p-value: *p < 0.05, ***p (C) Western blotting of *MYC* in Cas9-expressing MB157 cells transduced with control sgRNAs or sgRNAs simultaneously targeting *MYC* E1 and E5 enhancers as in (B). β-actin is loading control. (D) Representative histograms of CellTrace Violet (CTV) dilution in Cas9-expressing MB157 cells after transduction with control sgRNAs or sgRNAs simultaneously targeting *MYC* E1 and E5 enhancers as in (B). Unstained cells and freshly stained CTV cells are negative and positive controls, respectively. Barplot showing the average geometric mean fluorescent intensity (MFI) ± SD, n=3 biological replicates, representative of 3 independent experiments. t-test p-value: ***p < 1E-03. (E) Relative cell growth rates in Cas9-expressing MB157 cells after transduction with control sgRNAs or sgRNAs simultaneously targeting *MYC* E1 and E5 enhancers as in (B). Data represent mean ± SEM of n=8-10 biological replicates from 2 independent experiments. Day 9 data t-test p-value: ***p < 1E-03. (F) Notch-promoted loops (ΔL) linking Notch-activated enhancers (ΔE) to *CCND1* in MB157. Top panel: virtual 4C plot depicting the normalized interaction frequency from *CCND1* promoter viewpoint. ChIP-seq tracks showing Notch-sensitive NICD1 and RBPJ occupancy, and Notch-sensitive H3K27ac load marked with gray box. HiChIP arcs displaying normalized significant interactions between *CCND1* promoter and distal enhancers in Notch-on (top, DMSO) and Notch-off (bottom, GSI) (paired t-test p-value < 1E-15). Bottom track indicating *CCND1* Ensembl gene position. (G) Barplots showing qRT-PCR measurements of *CCND1* mRNA after transduction of Cas9-expressing MB157 cells with control sgRNAs, sgRNAs targeting the E1, E2 or both reveal independency of Notch-sensitive E1 and E2 *CCND1* enhancers. Data represent mean ± SEM of n=3-5 independent experiments. t-test p-value: **p < 0.01, ***p < 1E-03. (H) Western blotting of *CCND1* in Cas9-expressing MB157 cells transduced with control sgRNAs or sgRNAs simultaneously targeting *CCND1* E1 and E2 enhancers as in (G). Tubulin is loading control. (I) Representative histograms of CellTrace Violet (CTV) dilution in Cas9-expressing MB157 cells after transduction with control sgRNAs or sgRNAs simultaneously targeting *CCND1* E1 and E2 enhancers as in (G). Unstained cells and freshly stained CTV cells are negative and positive controls, respectively. Barplot showing the average geometric mean fluorescent intensity (MFI) ± SD, n=3 biological replicates, representative of 3 independent experiments. t-test p-value: ***p < 1E-03. (J) Representative flow plots of EdU incorporation in unstained cells, Cas9-expressing MB157 cells transduced with control sgRNAs or sgRNAs simultaneously targeting *CCND1* E1 and E2 enhancers as in (G). Barplot showing the average geometric mean fluorescent intensity (MFI) ± SD, n=3 biological replicates, representative of 2 independent experiments. t-test p-value: **p < 0.01.

We examined genes which were activated by Notch-promoted interactions to Notch-insensitive enhancers (Figure 3D mode b), to further analyze this previously unappreciated mode of Notch-dependent gene regulation. Virtual 4C (v4C) analysis of the TNBC-specific long noncoding RNA (lncRNA) *LINC00511*^63^ showed gain of contacts between the promoter and Notch-insensitive 3’ enhancers E3 (Figure 3I). Normalized contact tracks showed that in addition to *LINC00511*, Notch binding significantly increased the frequency of contacts between anchors linking Notch-bound Notch-insensitive enhancers and the Notch-sensitive gene *SOX9* (mode b, paired t-test p-value < 1E-03, Figure 3I). The same mode of Notch regulation of *LINC00511* and *SOX9* also operates in HCC1599 cells (model b, paired t-test p-value = 2E-03, Figure S3G). Together, these data are consistent with the ability of Notch to promote or strengthen certain enhancer-promoter interactions to activate direct Notch target genes, independent of changes in enhancer H3K27ac level.

In some instances, a common enhancer spatially co-regulates multiple genes through looping interactions with the promoters of each gene^57,64–68^. In MB157 and HCC1599 TNBC cells, Notch activation promotes looping interactions involved in spatial co-regulation of the kinase *RIPK4* and the serine protease *TMPRSS2* (Figures 3J and S3H), both of which are implicated in breast cancer pathogenicity^69–73^. Based on normalized contact tracks (Figures 3J and S3H), Notch activation significantly increased transcript abundance and contact frequency of *RIPK4* and *TMPRSS2* promoters to common Notch-bound and -activated enhancers, located 155 and 150 Kb away, respectively. Taken together, these results suggest that in addition to activating enhancers already in contact with promoters, Notch signaling can promote and strengthen physical interactions between promoters and enhancers in TNBC.

### Notch preferentially targets hyperconnected 3D regulatory cliques in TNBC

In addition to long-range enhancer-promoter loops, enhancer-enhancer and promoter-promoter interactions are implicated in gene control^38,74–79^. To examine the higher-order structure of regulatory interactions in Notch-mutated TNBC, we integrated our high-resolution connectivity maps and epigenomic data to annotate the regulatory loop anchors connecting enhancer or promoter elements (Table S7). We observed that a multiplicity of enhancer and promoter interactions were common in the Notch-mutated TNBC genomes, with each element on average connecting to 6 other regulatory elements (Figure S4A), as reported in other cell types^80,81^. Enhancer-promoter interactions accounted for only 30% of the long-range connections between regulatory elements in the Notch-mutated TNBC genomes (Figure 4A), also consistent with frequencies reported in other studies of mammalian cells^36^. Notably, only 30% of Notch-sensitive loops connected an enhancer to a promoter, while the majority linked pairs of enhancers (64%) (Figure 4A). In addition, 18% of interactions were between promoter pairs, in line with other reports suggesting the existence of regulatory promoter-promoter interactions^75,77^.

To globally model the higher-order structure of interactions involving Notch-sensitive regulatory interactions in TNBC cells, we used undirected graph mathematical abstraction^82^, and algorithmically^83^ searched for groups of densely connected enhancers and promoters with high intra-group and sparse inter-group interactions (see Method). We called these groups of highly interconnected elements “3D cliques”. We observed a significant asymmetry in the 3D clique connectivity distributions (Figures 4B and 4C, Table S8). Although 90% of 3D cliques contained less than 20 interactions, nearly 140 cliques were categorized as hyperconnected 3D cliques and had more than 100 interactions in either MB157 or HCC1599 (Figure 4B). The clique containing *MYC* was identified as a hyperconnected 3D clique, ranking among the top 10 most connected 3D cliques in both HCC1599 and MB157 cells (Figure 4B). Inspection of the *MYC* cliques in MB157 and HC1599 cells showed that Notch significantly promoted up to 46% of its constituent enhancers and more than 30% of interactions among and between its promoters and enhancers (Figures 4D and S4C).

This observation led us to ask whether Notch preferentially targets highly connected cliques in TNBC. We observed that the Notch-bound cliques exhibited significantly more connectivity than cliques lacking Notch binding (Wilcoxon rank sum p-value < 1E-15, Figures 4E and S4B). More connected cliques also contained significantly more Notch-sensitive loops (Wilcoxon rank sum p-value < 1E-15, Figure 4F). Furthermore, promoters connecting to Notch-activated enhancers through Notch-promoted loops interacted with more enhancers on average (Wilcoxon rank sum p-value < 1E-15, Figure S4D) and fell within cliques with higher connectivity (Wilcoxon rank sum p-value < 1E-02, Figures 4G and 4H). More importantly, the top 25% of the most connected cliques were enriched for direct Notch target genes relative to other cliques after correcting for clique connectivity (Wilcoxon rank sum p-value < 1E-03, Figures 4I and 4J). Direct Notch-activated genes within hyperconnected 3D cliques, such as *MYC* (Figure 4D and S4C), were associated with processes and pathways that have important functions in TNBC pathobiology (Figure S4E, Table S9). Overall, these results suggest that oncogenic Notch activates not only large stretches of enhancers as reported^17,20^, but also promotes regulatory DNA loops linking multiple distally located enhancers to their target genes.

### Perturbation of Notch-bound interacting enhancers reveals cooperativity in the *MYC* clique

Multiple enhancers are found in the several megabase region flanking the *MYC* gene body, but which transcription factors regulate *MYC* via these enhancers is not completely understood^18–21,84–89^. We observed that the *MYC* enhancers identified in Notch-mutated TNBC were also active in other TNBC lines but not non-TNBC cell lines (Figure S5A). Based on our observations that *MYC* enhancers in TNBC cells are organized into a hyperconnected 3D clique with frequent inter-enhancer interactions (Figures 4B, 4D, and S4C, Table S8), and that several distinct Notch-bound super-enhancers lie 5’ of the *MYC* promoter (Figures 4D and 5A), we next asked whether *MYC* enhancers cooperatively regulate *MYC* expression. Our data showed that among all *MYC* enhancer pairs (labeled E1 to E5 in Figure 5A), the strongest RBPJ/NOTCH1 ChIP-seq signals and the largest Notch-dependent changes in H3K27ac were observed in E1 and E5 (Figure 5A), located 451 and 65 Kb 5’ of the *MYC* promoter. Normalized contact tracks also showed that E1 and E5 enhancer pair extensively interacted (supported by 462 normalized HiChIP reads). Based on the relative magnitudes of the HiChIP signal in the Notch-on and Notch-off states, the E1-E5 interaction frequency was reduced by 8-fold after Notch inhibition, whereas the contact frequencies between the *MYC* promoter and its distal enhancers were attenuated by ~4-fold (Figure 5A).

To test the functional role of the E1 and E5 enhancers in cooperatively regulating *MYC*, we used CRISPR/Cas9 genome editing (Figure S5C) to mutate the consensus RBPJ binding motifs in E1 and E5 (Figures 4D, 5A and S5C). Mutation of RBPJ binding sites at E1 or E5 resulted in a 15% or 25% decrease in *MYC* expression, respectively; while simultaneous targeting yielded more than a 50% reduction in *MYC* transcript abundance (Figure 5B) and greatly reduced the MYC protein amount (Figure 5C). Dual targeting of these enhancers also suppressed cell proliferation as assessed by cell-trace violet staining (t-test p-value < 1E-03, Figure 5D) and cell counts (t-test p-value < 1E-03, Figure 5E). Overall, these data suggest that Notch transcription complexes increase *MYC* expression by promoting higher order complex interactions involving cooperating E1 and E5 enhancers and the *MYC* promoter.

### Notch-bound non-interacting enhancers independently regulate Cyclin D1 (*CCND1*)

Our data showed that Notch upregulates *CCND1* transcripts in TNBC, as reported^43,90^, but not in Notch-mutated MCL and T-ALL (Figure S3C). In MB157, the *CCND1* promoter and associated enhancers were organized into a 3D clique of moderate connectivity (46 interactions), which was substantially smaller than the *MYC* clique (682 interactions) (Figure S5D, Table S8). Analysis of the *CCND1* locus in MB157 cells demonstrated a 1.4-fold or greater reduction in contact frequency between the *CCND1* promoter and Notch-responsive enhancers after Notch inhibition (paired t-test p-value < 1E-09, Figures 5F and S5D). However, the enhancer-enhancer interaction frequency between the two strongest Notch-bound *CCND1* enhancers, E1 and E2, was 12-fold lower than the interactions between the *MYC* E1 and E5 enhancers (Figures 5A and 5F). To test for cooperativity between the *CCND1* E1 and E2 enhancers, we again used CRISPR/Cas9 targeting (Figures 5F, S5E, and S5F). Single targeting of RBPJ motifs in the E1 and E2 enhancers led to 55% and 35% decreases in *CCND1* expression, respectively (Figure 5G). However, simultaneous targeting of E1 and E2 did not show additive or cooperative effects on *CCND1* transcript abundance (Figure 5G). Nevertheless, we did observe significant effects of E1 and E2 dual targeting on CCND1 protein amount (Figure 5H), cell proliferation and cell cycle progression, as assessed by cell-trace violet staining (t-test p-value < 1E-03, Figure 5I) and EdU incorporation (t-test p-value < 1E-02, Figure 5J), respectively. Thus, like *MYC*, *CCND1* is another proto-oncogene that is dysregulated in TNBC by Notch through Notch-sensitive looping interactions involving lineage-specific distal enhancers. Furthermore, our data hint that enhancer-enhancer interactions could potentially influence the cooperativity between distal enhancers in transcription regulation.

### TNBC Notch regulatory modes are generalizable to Notch-mutated MCL

The observation that Notch preferentially promoted or strengthened regulatory loops in highly connected 3D cliques of Notch-mutated TNBC, led us to investigate whether the same relationships hold in other Notch-driven malignancies, such as MCL. We first analyzed the long-range interactions between two previously characterized Notch-activated enhancers located 525 Kb and 433 Kb 5’ of the *MYC* promoter in MCL Rec-1 cells^18^. The analysis of Rec-1 cohesin HiChIP showed that these two Notch-activated *MYC* enhancers (Figure S6A) interacted frequently (116.2 normalized reads) and that Notch inhibition significantly reduced this interaction (t-test p-value < 1E-15, Figure 6A). Based on these chromatin looping data and our analysis of Notch-activated *MYC* enhancers in TNBC, we conjectured that the two MCL-restricted Notch-activated enhancers cooperatively control *MYC* expression. This hypothesis was confirmed in our published work where use of CRISPR-Cas9-KRAB repressors showed that these two enhancers cooperate to regulate *MYC* in MCL Rec-1 cells^18^. To extend this analysis genome-wide, we first assessed the Notch-sensitivity of intradomain interactions and their relationship to active and repressed chromatin in Rec-1 cells. As in Notch-mutated TNBC cells (Figures 2D-G), Notch inhibition decreased the intradomain interaction frequencies of more than 130 contact domains by more than 4-fold (Figure 6B), and this reduction preferentially occurred in active chromatin domains (Wilcoxon rank sum p-value < 1E-15, Figures 6B and 6C). In Notch-mutated MCL Rec-1 cells, the integration of chromatin conformation, epigenomic, and transcriptomic data sets again showed that direct Notch target genes with the greatest increase in transcription were those with Notch-activated enhancers and Notch-promoted looping interactions (Wilcoxon rank sum p-value < 0.03, Figure 6D and Figure 3D mode a). In addition to *MYC*, *LYN*, a direct Notch target gene that is essential for B-cell receptor activity^18,91^, also showed Notch-promoted enhancer-promoter contacts (paired t-test p-value < 1E-15, Figure S6B and Figure 3D mode a). *SH2B2*, a gene coding for an adaptor protein with an important role in B-cell development and activation^92,93^, was regulated by interaction between Notch-insensitive enhancers and the *SH2B2* promoter through Notch-promoted loops (paired t-test p-value < 1E-15, Figure S6C and Figure 3D mode b). The inspection of HiChIP data from Rec-1 cells also identified Notch-promoted loops that permit spatial co-regulation of two genes from a shared Notch-activated enhancer (Figure S6D). Together, these data confirm that distinct Notch regulatory modes identified in TNBC also apply to Notch-mutated MCL.

**Figure 6.**
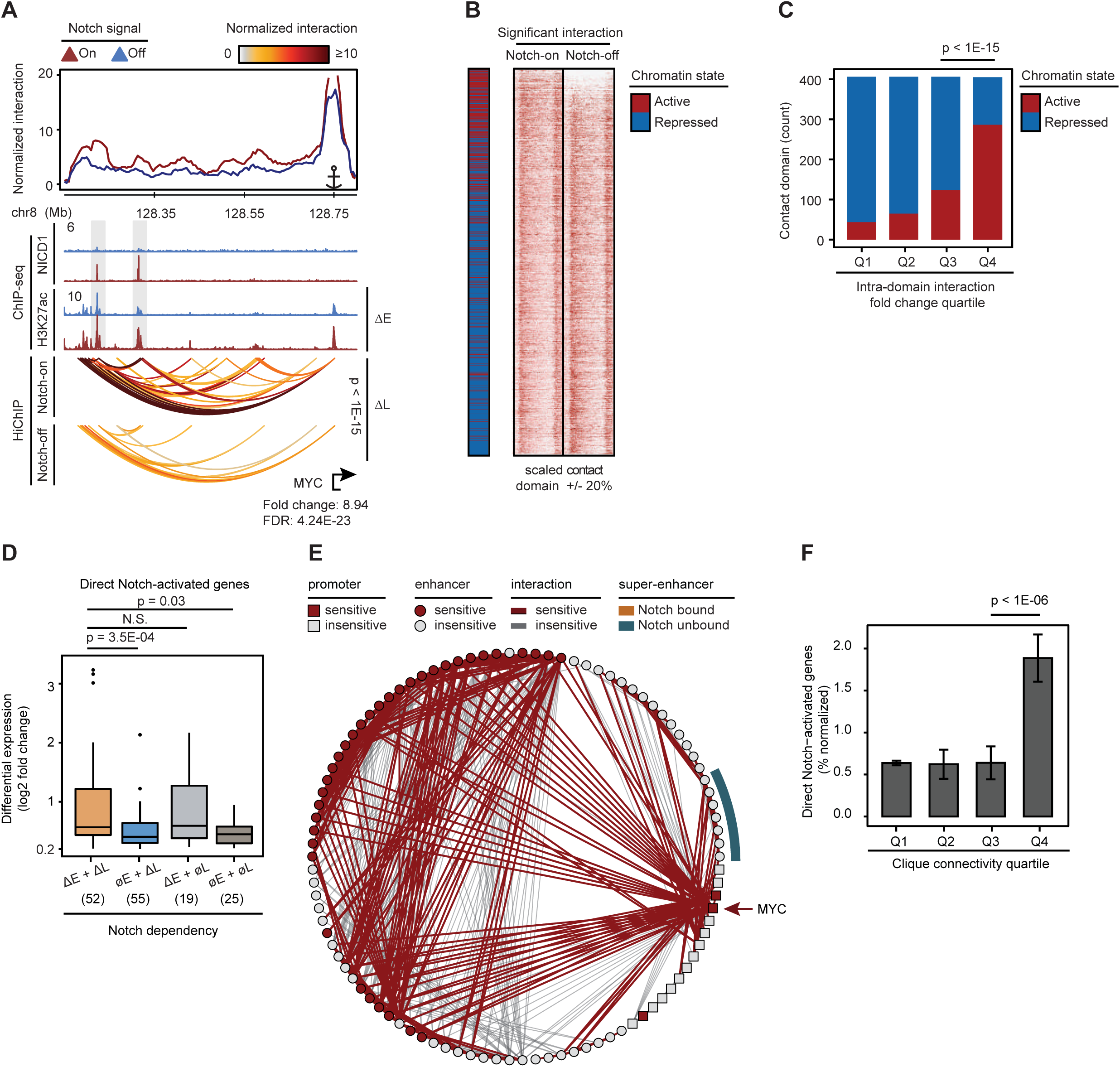
Activation of MCL Notch direct target genes by Notch-promoted and preformed enhancer-promoter contacts. (A) Notch-promoted loops (ΔL) linking Notch-activated enhancers (ΔE) to *MYC* in MCL Rec-1 cells. Top panel: virtual 4C plot depicting the normalized interaction frequency from *MYC* promoter viewpoint. ChIP-seq tracks showing Notch-sensitive NICD1 and RBPJ occupancy, and Notch-sensitive H3K27ac load marked with gray box. HiChIP arcs displaying normalized significant interactions among *MYC* promoter and distal enhancers in Notch-on (top, DMSO) and Notch-off (bottom, GSI) (paired t-test p-value < 1E-15). Bottom track indicating *MYC* Ensembl gene position and its expression fold change and FDR as determined by DESeq2. (B) Heatmap of normalized significant interactions at scaled and flanked contact domains in Rec-1 in Notch-on (DMSO) and Notch-off (GSI) conditions. Contact domains are ranked by change (log2 fold change) of total intradomain contacts with the chromatin states indicated on the left. The overall differential intradomain contact frequency is significant (paired t-test p-value < 1E-15). (C) Barplot depicting the number of active/repressed contact domains per quartile of total intradomain contact frequency change in Rec-1 (proportion test p-value < 1E-15). (D) Boxplot showing differential (log_2_ fold change) gene expression of direct Notch-activated genes in Rec-1 categorized by Notch-dependency of interacting Notch-bound enhancers and loops. Notch-bound and -promoted loops (ΔL), Notch-bound and -activated enhancers (ΔE), Notch-bound but Notch-insensitive loops (ØL), and Notch-bound but Notch-insensitive enhancers (ØE). (E) Circos plot showing the clique associated with *MYC* in Rec-1. Red-marked circle (square) and line depicting Notch-sensitive enhancer (promoter) and significant long-range interactions, respectively. (F) Barplot depicting the average ± SEM corrected percentage of direct Notch-activated genes per quartile of clique total connectivity distribution in Rec-1 (Wilcoxon rank sum p-value < 1E-06).

Analysis of HiChIP data revealed that the Rec-1 genome is also organized into 3D regulatory cliques consisting of densely interconnected enhancers and promoters. A significant asymmetry was also observed in the Rec-1 clique connectivity distribution (Figure S6E). Cliques with higher connectivity were enriched for direct Notch target genes, including *MYC* and genes involved in the B-cell signaling response and regulation (e.g. *IL10RA*, *PAX5*, *CR2*) (Figures 6E, S6E, and S6F). Further assessment of Rec-1 cliques showed significant enrichment for direct Notch target genes in highly connected 3D cliques with Notch-sensitive enhancers and looping interactions (Wilcoxon rank sum p-value < 1E-06, Figures 6E and 6F). These direct Notch-activated genes were associated with processes and pathways with known roles in B cell biology and lymphomagenesis (Figure S6G, Table S10). Overall, these results suggest that Notch signaling controls not only MCL transcriptional enhancers, as reported^18^, but also strengthens or promotes enhancer-promoter looping interactions in MCL cliques to regulate critical B cell pathways.

### Notch reactivation rescues regulatory looping interactions

Our data revealed that Notch inhibition “decommissions” regulatory loops, leading to down-regulation of Notch target genes. If Notch-mediated regulatory loops are dynamically regulated by Notch, these loops should be rapidly restored following Notch reactivation by GSI washout (Figure 7A). Analysis of cohesin HiChIP following GSI-washout (Figure S7A), as expected, showed no change in contact domains with Notch reactivation (Figures S7B and S7C). However, long-range regulatory interactions, including enhancer-enhancer, enhancer-promoter and promoter-promoter interactions, were restored after Notch reactivation (paired t-test p-value < 1E-15, Figure 7B). Specifically, Notch inhibition significantly decreased 412 interactions between Notch direct target genes and their enhancers (fold change > 1.4, FDR < 1E-10), of which 74% were completely recovered following Notch reactivation by GSI washout (paired t-test p-value < 1E-15, Figure 7C), including looping interactions involving *MYC* and *CCND1* (Figures 7D and S7D). Furthermore, Notch reactivation recovered 74% of the Notch-dependent chromatin loops in the *MYC* clique (Figure 7E). Together, these results support a model in which loading of Notch transcription complexes onto regulatory elements has widespread effects on looping interactions involving the genomes of Notch-mutated cancer cells.

**Figure 7.**
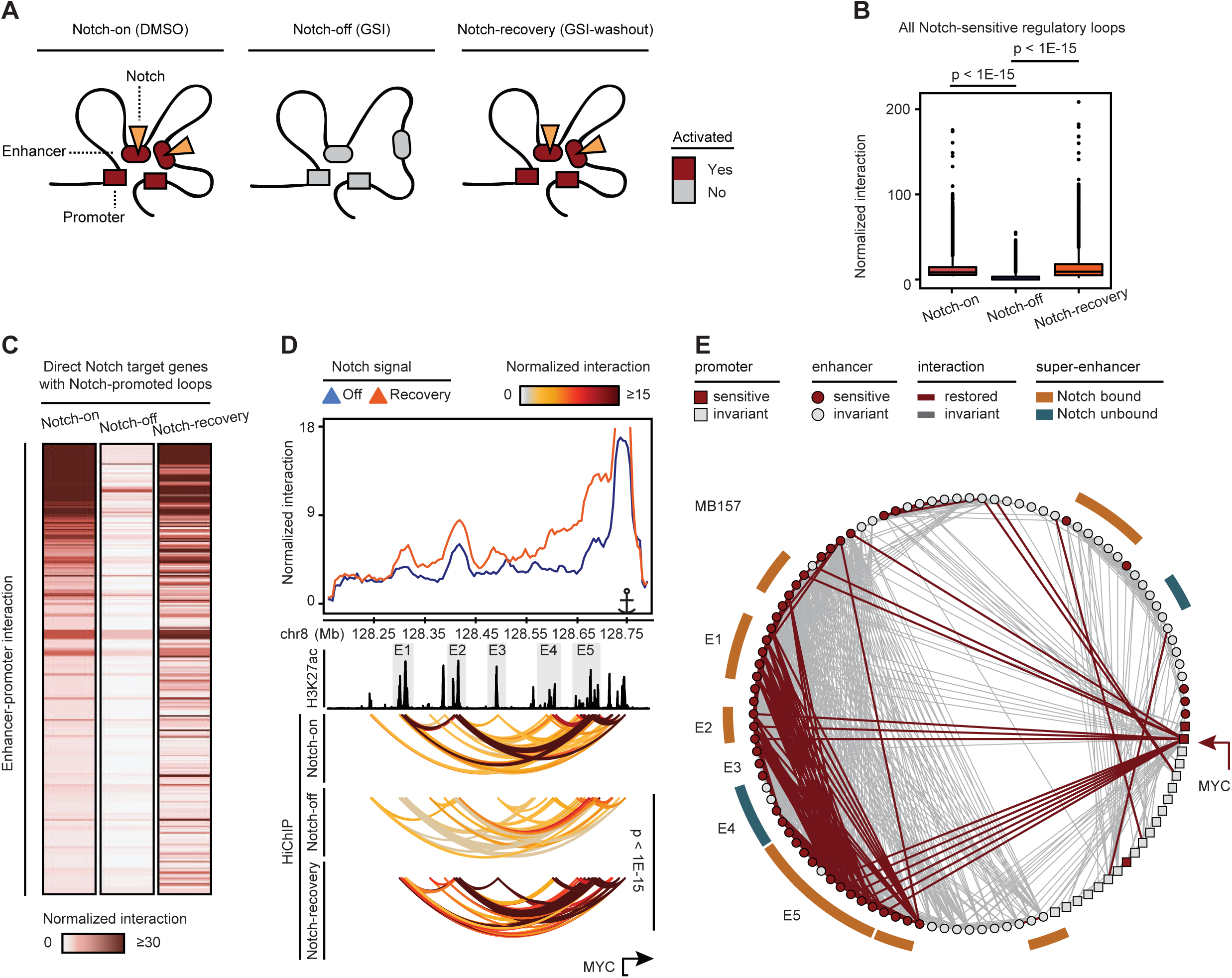
Notch reactivation rescues regulatory interactions. (A) Scheme showing reduced transcriptional activities as a result of loop and enhancer deactivation upon Notch-inhibition by GSI and restoration after Notch recovery by GSI-washout. (B) Boxplots displaying the normalized interaction frequency of Notch-promoted enhancer-enhancer, enhancer-promoter, and promoter-promoter interactions in Notch-on (DMSO), Notch-off (GSI) and Notch-recovery (GSI-washout) conditions in MB157 (paired t-test p-value < 1E-15). (C) Heatmap showing the normalized enhancer-promoter interactions of direct Notch targets with Notch-promoted loops (ΔL) in Notch-on (DMSO), Notch-off (GSI) and Notch-recovery (GSI-washout) conditions. Each row is a pair of enhancer-promoter interaction sorted in descending order of Notch differential effect (log_2_ fold change). The overall differences between enhancer-promoter interactions in Notch-on versus Notch-off and Notch-recovery versus Notch-off are significant (paired t-test p-value < 1E-15). (D) Notch recovery rescues interactions at *MYC* locus in MB157. Top panel: virtual 4C plot depicting the normalized interaction frequency from *MYC* promoter viewpoint. ChIP-seq tracks showing H3K27ac load. *MYC* enhancers are marked with gray box as in Figure 5. HiChIP arcs displaying normalized significant interactions between *MYC* promoter and distal enhancers in Notch-on (top, DMSO), Notch-off (middle, GSI), and Notch-recovery (bottom, GSI-washout). (E) Interactions of the *MYC* clique in MB157 were recovered upon Notch reactivation. Red lines represent interactions significantly decreased in Notch inhibition and restored in Notch-recovery (fold change > 1.4, enrichR FDR < 0.05).

## Discussion

Chromatin architecture dynamics in response to oncogenic transcription factors are not well understood. Here, we used the response to oncogenic subversion of the developmental transcription factor Notch in TNBC and MCL, two cancers with frequent Notch-activating mutations, to examine the impact of aberrant transcription factor activity on long-range regulatory loops in tumors. Our data corroborate earlier studies showing that Notch binding events often associate with increased histone acetylation, not necessarily in proximity to transcribed genes^17,18^. We analyzed the impact of Notch on chromatin state and conformation and the consequence of such changes on transcriptional outputs. Our high-resolution chromatin conformation maps of Notch-mutated tumors revealed that oncogenic Notch signaling differentially affects the 3D genome organization hierarchy. While chromatin contact domains are largely independent of Notch-responsive transcription factor and conserved across Notch-mutated cancer cells, we strikingly find that in addition to activating enhancers already in contact with genes, oncogenic Notch leads to a gain in contact frequency between activated genes and distal Notch-bound enhancers. Based on our data, we propose that Notch relies on four distinct regulatory modes, defined by the combination of enhancer and/or loop acquisition, to control its direct target genes. Importantly, concomitant Notch-mediated enhancer activation and gain in enhancer-promoter contact frequency leads to a larger increase in the expression of direct Notch target genes. Together, our data suggest that in addition to activating enhancers already in contact with transcriptional targets, oncogenic transcription factors may promote or stabilize regulatory interactions between promoter and enhancers to activate transcription.

The study of long-range regulation of gene expression by signal-dependent transcription factors resulted in conflicting results on whether transcription factors join existing chromatin loops or remodel the loops themselves to activate gene expression^4,94–97^. Discrepancies between studies on transcription factor-mediated long-range DNA loop changes may be due to differences in resolution, methodology, or the nature of the given transcription factor. Recent studies showed that the lineage-specific chromatin structure is established in tissue progenitor cells and is further remodeled in terminal differentiation^98,99^. Here, we demonstrate that toggling between active and inactive Notch signaling in cancer cells controls gene expression by reorganizing long-range regulatory interactions. This observation has implications for targeting undruggable proto-oncogenes with long-range looping interactions in Notch-dependent tumors.

Notch transcription complexes recruit other transcriptional regulators, such as chromatin enzymes^13^ and transcriptional coactivators^16^. Our results suggest that Notch-binding is selectively required for enhancer activation and promoting contacts among and between enhancers and promoters, but Notch transcription complexes binding is not sufficient to determine regulatory loops and chromatin state dynamics. The exact mechanism explaining various Notch regulatory modes and the specificity requirements of Notch-promoted regulatory DNA loops remains to be determined. Notch could potentially modify regulatory loops intrinsically (for example, by dimerization^58,100,101^) or interact with known architectural proteins^2,102^. Further studies are needed to determine the impact of cooperation between cognate transcription factor binding and chromatin remodelers, among other factors.

Chromatin organization is a major determinant of regulatory interactions. By definition, interactions between regulatory DNA elements are more likely to occur within contact domains than across them^26,30,31,55^. Nevertheless, sequences in different domains of a chromosome interact, albeit at a much lower frequency, and may be important for proper gene control. Our high-resolution regulatory connectivity maps identified complexities of localized and long-range enhancer and promoter sharing. We identified spatially interacting communities of regulatory elements, termed 3D cliques, independent of their contact domains. By systematically delineating clusters of frequently interacting enhancers and promoters in the regulatory interaction graph (i.e. 3D regulome) of Notch-mutated tumors, we expanded on previous observations of pairwise contacts between super-enhancers, high interactions among constitutive elements of super-enhancer regions, and promiscuous locally interacting regions^103–105^. Here, we show that long-range regulatory loops of Notch-mutated cancer cells coalesce enhancers and promoters to form 3D regulatory cliques. Oncogenic Notch preferentially promotes enhancers and DNA loops in hyperconnected 3D cliques. To this end, oncogenic Notch not only activates large stretches of enhancers (or super-enhancers) in the 1D genome as reported in Notch-dependent T cell leukemia^17,21^, it also induces long-range regulatory interactions among multiple distal enhancers, including distinct super-enhancers, to promote activity of *MYC* (Figures 4D and S4C). These findings suggest that the entire regulatory interaction map should be taken into consideration for enhancer editing, as cooperativity between enhancers to control gene expression may not solely depend on individual promoter interaction, but could depend on other factors such as connectivity among enhancers.

Our data also suggest that by targeting hyperconnected 3D cliques of key oncogenes such as *MYC*, Notch uses a multiplicity of distal enhancers and enhancer-enhancer interactions to maximize Notch-driven pathogenic transcription outputs. These observations suggest that while the genomic loci with a high frequency of chromatin interactions are highly enriched for super-enhancers^103,105^, several super-enhancers could distally interact to control key oncogenes. Notch activation of large cliques formed by pre-existing and gained loops is reminiscent of “active chromatin hub” formation at the beta-globin locus in which multiple distal sites loop to the active beta-globin genes during specific stages of erythrocyte development^106–108^. It is possible to speculate that the formation of larger aggregates of regulatory elements into 3D cliques might increase the concentration of transcription coactivators to form phase-separated condensates at spatial aggregates of super-enhancers that compartmentalize and concentrate the transcription apparatus ^65,109,110^. Hence, the observation of exceptionally large interacting cliques of regulatory elements advocates in favor of the applicability of the nuclear condensate model for the Notch-mediated activation of key cancer cell genes. Together, our results implicate reorganization of regulatory loops as an instructive factor for implementing oncogenic transcription factors-driven transcriptional programs.

## Acknowledgements

We thank Howard Chang for providing HiChIP protocol. This work was supported by Abramson Cancer Center Cooper Award, Abramson Family Cancer Research Institute Investigator Award, and Translational Medicine and Therapeutics program for Transdisciplinary Awards Program in Translational Medicine and Therapeutics (to R.B.F.), Penn Epigenetics Institute Pilot Award (to R.B.F. and W.S.P.), R01-CA-215518 (to W.S.P.), T32-CA009140 (to G.W.S.), and LLS-5456-17 (to J.P.)

## Authors Contributions

Conceptualization: R.B.F., J.P., Y.Z.; Methodology: R.B.F., W.S.P., J.P., Y.Z., M.F., G.V., E.F.J.; Investigation: R.B.F., W.S.P., J.P., Y.Z.; Formal Analysis: R.B.F., J.P., Y.Z., M.F., N.G, G.W.S., S.C.N., K.S.R., Y.S., Z.Z., S.C.L.; Resources and Reagents: R.B.F., W.S.P., E.F.J., J.S., S.C.L., M.J.K., S.C.B., M.R.M., J.C.A., G.V.; Writing-Original Draft: R.B.F., W.S.P., J.P., Y.Z.; Supervision: R.B.F., W.S.P.; Funding Acquisition: R.B.F., W.S.P.

## References

1 Dixon, J. R. et al. Topological domains in mammalian genomes identified by analysis of chromatin interactions. Nature 485, 376–380, doi:10.1038/nature11082 (2012).

2 Hnisz, D., Day, D. S. & Young, R. A. Insulated Neighborhoods: Structural and Functional Units of Mammalian Gene Control. Cell 167, 1188–1200, doi:10.1016/j.cell.2016.10.024 (2016).

3 Misteli, T. Higher-order genome organization in human disease. Cold Spring Harb Perspect Biol 2, a000794, doi:10.1101/cshperspect.a000794 (2010).

4 Jin, F. et al. A high-resolution map of the three-dimensional chromatin interactome in human cells. Nature 503, 290–294, doi:10.1038/nature12644 (2013).

5 Flavahan, W. A. et al. Insulator dysfunction and oncogene activation in IDH mutant gliomas. Nature 529, 110–114, doi:10.1038/nature16490 (2016).

6 Hnisz, D. et al. Activation of proto-oncogenes by disruption of chromosome neighborhoods. Science 351, 1454–1458, doi:10.1126/science.aad9024 (2016).

7 Katainen, R. et al. CTCF/cohesin-binding sites are frequently mutated in cancer. Nat Genet 47, 818–821, doi:10.1038/ng.3335 (2015).

8 Lawrence, M. S. et al. Discovery and saturation analysis of cancer genes across 21 tumour types. Nature 505, 495–501, doi:10.1038/nature12912 (2014).

9 Taberlay, P. C. et al. Three-dimensional disorganization of the cancer genome occurs coincident with long-range genetic and epigenetic alterations. Genome Res 26, 719–731, doi:10.1101/gr.201517.115 (2016).

10 Wu, P. et al. 3D genome of multiple myeloma reveals spatial genome disorganization associated with copy number variations. Nat Commun 8, 1937, doi:10.1038/s41467-017-01793-w (2017).

11 Bray, S. J. Notch signalling in context. Nat Rev Mol Cell Biol 17, 722–735, doi:10.1038/nrm.2016.94 (2016).

12 Aster, J. C., Pear, W. S. & Blacklow, S. C. The Varied Roles of Notch in Cancer. Annu Rev Pathol 12, 245–275, doi:10.1146/annurev-pathol-052016-100127 (2017).

13 Oswald, F. et al. p300 acts as a transcriptional coactivator for mammalian Notch-1. Mol Cell Biol 21, 7761–7774, doi:10.1128/MCB.21.22.7761-7774.2001 (2001).

14 Mulligan, P. et al. A SIRT1-LSD1 corepressor complex regulates Notch target gene expression and development. Mol Cell 42, 689–699, doi:10.1016/j.molcel.2011.04.020 (2011).

15 Yatim, A. et al. NOTCH1 nuclear interactome reveals key regulators of its transcriptional activity and oncogenic function. Mol Cell 48, 445–458, doi:10.1016/j.molcel.2012.08.022 (2012).

16 Fryer, C. J., White, J. B. & Jones, K. A. Mastermind recruits CycC:CDK8 to phosphorylate the Notch ICD and coordinate activation with turnover. Mol Cell 16, 509–520, doi:10.1016/j.molcel.2004.10.014 (2004).

17 Wang, H. et al. NOTCH1-RBPJ complexes drive target gene expression through dynamic interactions with superenhancers. Proc Natl Acad Sci U S A 111, 705–710, doi:10.1073/pnas.1315023111 (2014).

18 Ryan, R. J. H. et al. A B Cell Regulome Links Notch to Downstream Oncogenic Pathways in Small B Cell Lymphomas. Cell Rep 21, 784–797, doi:10.1016/j.celrep.2017.09.066 (2017).

19 Fabbri, G. et al. Common nonmutational NOTCH1 activation in chronic lymphocytic leukemia. Proc Natl Acad Sci U S A 114, E2911–E2919, doi:10.1073/pnas.1702564114 (2017).

20 Yashiro-Ohtani, Y. et al. Long-range enhancer activity determines Myc sensitivity to Notch inhibitors in T cell leukemia. Proc Natl Acad Sci U S A 111, E4946–4953, doi:10.1073/pnas.1407079111 (2014).

21 Herranz, D. et al. A NOTCH1-driven MYC enhancer promotes T cell development, transformation and acute lymphoblastic leukemia. Nat Med 20, 1130–1137, doi:10.1038/nm.3665 (2014).

22 Hu, Z. & Tee, W. W. Enhancers and chromatin structures: regulatory hubs in gene expression and diseases. Biosci Rep 37, doi:10.1042/BSR20160183 (2017).

23 Spitz, F. Gene regulation at a distance: From remote enhancers to 3D regulatory ensembles. Semin Cell Dev Biol 57, 57–67, doi:10.1016/j.semcdb.2016.06.017 (2016).

24 de Laat, W. & Duboule, D. Topology of mammalian developmental enhancers and their regulatory landscapes. Nature 502, 499–506, doi:10.1038/nature12753 (2013).

25 Visel, A., Rubin, E. M. & Pennacchio, L. A. Genomic views of distant-acting enhancers. Nature 461, 199–205, doi:10.1038/nature08451 (2009).

26 Kagey, M. H. et al. Mediator and cohesin connect gene expression and chromatin architecture. Nature 467, 430–435, doi:10.1038/nature09380 (2010).

27 Schmidt, D. et al. A CTCF-independent role for cohesin in tissue-specific transcription. Genome Res 20, 578–588, doi:10.1101/gr.100479.109 (2010).

28 Phillips-Cremins, J. E. et al. Architectural protein subclasses shape 3D organization of genomes during lineage commitment. Cell 153, 1281–1295, doi:10.1016/j.cell.2013.04.053 (2013).

29 Watrin, E., Kaiser, F. J. & Wendt, K. S. Gene regulation and chromatin organization: relevance of cohesin mutations to human disease. Curr Opin Genet Dev 37, 59–66, doi:10.1016/j.gde.2015.12.004 (2016).

30 Dowen, J. M. et al. Control of cell identity genes occurs in insulated neighborhoods in mammalian chromosomes. Cell 159, 374–387, doi:10.1016/j.cell.2014.09.030 (2014).

31 Lupianez, D. G. et al. Disruptions of topological chromatin domains cause pathogenic rewiring of gene-enhancer interactions. Cell 161, 1012–1025, doi:10.1016/j.cell.2015.04.004 (2015).

32 Nora, E. P. et al. Spatial partitioning of the regulatory landscape of the X-inactivation centre. Nature 485, 381–385, doi:10.1038/nature11049 (2012).

33 Rao, S. S. et al. A 3D map of the human genome at kilobase resolution reveals principles of chromatin looping. Cell 159, 1665–1680, doi:10.1016/j.cell.2014.11.021 (2014).

34 Tang, Z. et al. CTCF-Mediated Human 3D Genome Architecture Reveals Chromatin Topology for Transcription. Cell 163, 1611–1627, doi:10.1016/j.cell.2015.11.024 (2015).

35 Ji, X. et al. 3D Chromosome Regulatory Landscape of Human Pluripotent Cells. Cell Stem Cell 18, 262–275, doi:10.1016/j.stem.2015.11.007 (2016).

36 Weintraub, A. S. et al. YY1 Is a Structural Regulator of Enhancer-Promoter Loops. Cell 171, 1573–1588 e1528, doi:10.1016/j.cell.2017.11.008 (2017).

37 Mumbach, M. R. et al. HiChIP: efficient and sensitive analysis of protein-directed genome architecture. Nat Methods 13, 919–922, doi:10.1038/nmeth.3999 (2016).

38 Mumbach, M. R. et al. Enhancer connectome in primary human cells identifies target genes of disease-associated DNA elements. Nat Genet 49, 1602–1612, doi:10.1038/ng.3963 (2017).

39 Rowley, M. J. et al. Evolutionarily Conserved Principles Predict 3D Chromatin Organization. Mol Cell 67, 837–852 e837, doi:10.1016/j.molcel.2017.07.022 (2017).

40 Fang, R. et al. Mapping of long-range chromatin interactions by proximity ligation-assisted ChIP-seq. Cell Res 26, 1345–1348, doi:10.1038/cr.2016.137 (2016).

41 Crane, E. et al. Condensin-driven remodelling of X chromosome topology during dosage compensation. Nature 523, 240–244, doi:10.1038/nature14450 (2015).

42 Henkel, T., Ling, P. D., Hayward, S. D. & Peterson, M. G. Mediation of Epstein-Barr virus EBNA2 transactivation by recombination signal-binding protein J kappa. Science 265, 92–95 (1994).

43 Choy, L. et al. Constitutive NOTCH3 Signaling Promotes the Growth of Basal Breast Cancers. Cancer Res 77, 1439–1452, doi:10.1158/0008-5472.CAN-16-1022 (2017).

44 Stoeck, A. et al. Discovery of biomarkers predictive of GSI response in triple-negative breast cancer and adenoid cystic carcinoma. Cancer Discov 4, 1154–1167, doi:10.1158/2159-8290.CD-13-0830 (2014).

45 Wang, K. et al. PEST domain mutations in Notch receptors comprise an oncogenic driver segment in triple-negative breast cancer sensitive to a gamma-secretase inhibitor. Clin Cancer Res 21, 1487–1496, doi:10.1158/1078-0432.CCR-14-1348 (2015).

46 Robinson, D. R. et al. Functionally recurrent rearrangements of the MAST kinase and Notch gene families in breast cancer. Nat Med 17, 1646–1651, doi:10.1038/nm.2580 (2011).

47 Turner, N. et al. Integrative molecular profiling of triple negative breast cancers identifies amplicon drivers and potential therapeutic targets. Oncogene 29, 2013–2023, doi:10.1038/onc.2009.489 (2010).

48 Wendt, K. S. et al. Cohesin mediates transcriptional insulation by CCCTC-binding factor. Nature 451, 796–801, doi:10.1038/nature06634 (2008).

49 Narendra, V. et al. CTCF establishes discrete functional chromatin domains at the Hox clusters during differentiation. Science 347, 1017–1021, doi:10.1126/science.1262088 (2015).

50 Sofueva, S. et al. Cohesin-mediated interactions organize chromosomal domain architecture. EMBO J 32, 3119–3129, doi:10.1038/emboj.2013.237 (2013).

51 Dixon, J. R. et al. Chromatin architecture reorganization during stem cell differentiation. Nature 518, 331–336, doi:10.1038/nature14222 (2015).

52 Weng, A. P. et al. c-Myc is an important direct target of Notch1 in T-cell acute lymphoblastic leukemia/lymphoma. Genes Dev 20, 2096–2109, doi:10.1101/gad.1450406 (2006).

53 Bailis, W., Yashiro-Ohtani, Y. & Pear, W. S. Identifying direct Notch transcriptional targets using the GSI-washout assay. Methods Mol Biol 1187, 247–254, doi:10.1007/978-1-4939-1139-4_19 (2014).

54 Boettiger, A. N. et al. Super-resolution imaging reveals distinct chromatin folding for different epigenetic states. Nature 529, 418–422, doi:10.1038/nature16496 (2016).

55 Dowen, J. M. et al. Multiple structural maintenance of chromosome complexes at transcriptional regulatory elements. Stem Cell Reports 1, 371–378, doi:10.1016/j.stemcr.2013.09.002 (2013).

56 Phanstiel, D. H., Boyle, A. P., Heidari, N. & Snyder, M. P. Mango: a bias-correcting ChIA-PET analysis pipeline. Bioinformatics 31, 3092–3098, doi:10.1093/bioinformatics/btv336 (2015).

57 Ghavi-Helm, Y. et al. Enhancer loops appear stable during development and are associated with paused polymerase. Nature 512, 96–100, doi:10.1038/nature13417 (2014).

58 Deng, W. et al. Controlling long-range genomic interactions at a native locus by targeted tethering of a looping factor. Cell 149, 1233–1244, doi:10.1016/j.cell.2012.03.051 (2012).

59 Abravanel, D. L. et al. Notch promotes recurrence of dormant tumor cells following HER2/neu-targeted therapy. J Clin Invest 125, 2484–2496, doi:10.1172/JCI74883 (2015).

60 Cohen, B. et al. Cyclin D1 is a direct target of JAG1-mediated Notch signaling in breast cancer. Breast Cancer Res Treat 123, 113–124, doi:10.1007/s10549-009-0621-9 (2010).

61 Regan, J. L. et al. c-Kit is required for growth and survival of the cells of origin of Brca1-mutation-associated breast cancer. Oncogene 31, 869–883, doi:10.1038/onc.2011.289 (2012).

62 Kashiwagi, S. et al. c-Kit expression as a prognostic molecular marker in patients with basal-like breast cancer. Br J Surg 100, 490–496, doi:10.1002/bjs.9021 (2013).

63 Xu, S., Kong, D., Chen, Q., Ping, Y. & Pang, D. Oncogenic long noncoding RNA landscape in breast cancer. Mol Cancer 16, 129, doi:10.1186/s12943-017-0696-6 (2017).

64 Fanucchi, S., Shibayama, Y., Burd, S., Weinberg, M. S. & Mhlanga, M. M. Chromosomal contact permits transcription between coregulated genes. Cell 155, 606–620, doi:10.1016/j.cell.2013.09.051 (2013).

65 Fukaya, T., Lim, B. & Levine, M. Enhancer Control of Transcriptional Bursting. Cell 166, 358–368, doi:10.1016/j.cell.2016.05.025 (2016).

66 Sanborn, A. L. et al. Chromatin extrusion explains key features of loop and domain formation in wild-type and engineered genomes. Proc Natl Acad Sci U S A 112, E6456–6465, doi:10.1073/pnas.1518552112 (2015).

67 Schoenfelder, S. et al. Preferential associations between co-regulated genes reveal a transcriptional interactome in erythroid cells. Nat Genet 42, 53–61, doi:10.1038/ng.496 (2010).

68 Zhang, Y. et al. Chromatin connectivity maps reveal dynamic promoter-enhancer long-range associations. Nature 504, 306–310, doi:10.1038/nature12716 (2013).

69 Huang, X. et al. Phosphorylation of Dishevelled by protein kinase RIPK4 regulates Wnt signaling. Science 339, 1441–1445, doi:10.1126/science.1232253 (2013).

70 Liu, D. Q. et al. Increased RIPK4 expression is associated with progression and poor prognosis in cervical squamous cell carcinoma patients. Sci Rep 5, 11955, doi:10.1038/srep11955 (2015).

71 Luostari, K. et al. Type II transmembrane serine protease gene variants associate with breast cancer. PLoS One 9, e102519, doi:10.1371/journal.pone.0102519 (2014).

72 Partanen, J. I. et al. Tumor suppressor function of Liver kinase B1 (Lkb1) is linked to regulation of epithelial integrity. Proc Natl Acad Sci U S A 109, E388–397, doi:10.1073/pnas.1120421109 (2012).

73 Xing, P. et al. Clinical and biological significance of hepsin overexpression in breast cancer. J Investig Med 59, 803–810, doi:10.2310/JIM.0b013e31821451a1 (2011).

74 Hah, N., Murakami, S., Nagari, A., Danko, C. G. & Kraus, W. L. Enhancer transcripts mark active estrogen receptor binding sites. Genome Res 23, 1210–1223, doi:10.1101/gr.152306.112 (2013).

75 Li, G. et al. Extensive promoter-centered chromatin interactions provide a topological basis for transcription regulation. Cell 148, 84–98, doi:10.1016/j.cell.2011.12.014 (2012).

76 Melo, C. A. et al. eRNAs are required for p53-dependent enhancer activity and gene transcription. Mol Cell 49, 524–535, doi:10.1016/j.molcel.2012.11.021 (2013).

77 Mifsud, B. et al. Mapping long-range promoter contacts in human cells with high-resolution capture Hi-C. Nat Genet 47, 598–606, doi:10.1038/ng.3286 (2015).

78 Kim, T. K. et al. Widespread transcription at neuronal activity-regulated enhancers. Nature 465, 182–187 (2010).

79 Wang, K. C. et al. A long noncoding RNA maintains active chromatin to coordinate homeotic gene expression. Nature 472, 120–124, doi:10.1038/nature09819 (2011).

80 Marinic, M., Aktas, T., Ruf, S. & Spitz, F. An integrated holo-enhancer unit defines tissue and gene specificity of the Fgf8 regulatory landscape. Dev Cell 24, 530–542, doi:10.1016/j.devcel.2013.01.025 (2013).

81 Pope, B. D. et al. Topologically associating domains are stable units of replication-timing regulation. Nature 515, 402–405, doi:10.1038/nature13986 (2014).

82 Diestel, R. Graph theory. (Springer, 1997).

83 Blondel, V. D., Guillaume, J. L., Lambiotte, R. & Lefebvre, E. Fast unfolding of communities in large networks. J Stat Mech-Theory E, doi:Artn P10008 10.1088/1742-5468/2008/10/P10008 (2008).

84 Ahmadiyeh, N. et al. 8q24 prostate, breast, and colon cancer risk loci show tissue-specific long-range interaction with MYC. Proc Natl Acad Sci U S A 107, 9742–9746, doi:10.1073/pnas.0910668107 (2010).

85 Lin, C. Y. et al. Active medulloblastoma enhancers reveal subgroup-specific cellular origins. Nature 530, 57–62, doi:10.1038/nature16546 (2016).

86 Schuijers, J. et al. Transcriptional Dysregulation of MYC Reveals Common Enhancer-Docking Mechanism. Cell Rep 23, 349–360, doi:10.1016/j.celrep.2018.03.056 (2018).

87 Shi, J. et al. Role of SWI/SNF in acute leukemia maintenance and enhancer-mediated Myc regulation. Genes Dev 27, 2648–2662, doi:10.1101/gad.232710.113 (2013).

88 Sur, I. K. et al. Mice lacking a Myc enhancer that includes human SNP rs6983267 are resistant to intestinal tumors. Science 338, 1360–1363, doi:10.1126/science.1228606 (2012).

89 Wasserman, N. F., Aneas, I. & Nobrega, M. A. An 8q24 gene desert variant associated with prostate cancer risk confers differential in vivo activity to a MYC enhancer. Genome Res 20, 1191–1197, doi:10.1101/gr.105361.110 (2010).

90 Ronchini, C. & Capobianco, A. J. Induction of cyclin D1 transcription and CDK2 activity by Notch(ic): implication for cell cycle disruption in transformation by Notch(ic). Mol Cell Biol 21, 5925–5934 (2001).

91 Gauld, S. B. & Cambier, J. C. Src-family kinases in B-cell development and signaling. Oncogene 23, 8001–8006, doi:10.1038/sj.onc.1208075 (2004).

92 Iseki, M. et al. APS, an adaptor molecule containing PH and SH2 domains, has a negative regulatory role in B cell proliferation. Biochem Biophys Res Commun 330, 1005–1013, doi:10.1016/j.bbrc.2005.03.073 (2005).

93 Naudin, C., Chevalier, C. & Roche, S. The role of small adaptor proteins in the control of oncogenic signalingr driven by tyrosine kinases in human cancer. Oncotarget 7, 11033–11055, doi:10.18632/oncotarget.6929 (2016).

94 Kuznetsova, T. et al. Glucocorticoid receptor and nuclear factor kappa-b affect three-dimensional chromatin organization. Genome Biol 16, 264, doi:10.1186/s13059-015-0832-9 (2015).

95 Qiao, Y. et al. AP-1-mediated chromatin looping regulates ZEB2 transcription: new insights into TNFalpha-induced epithelial-mesenchymal transition in triple-negative breast cancer. Oncotarget 6, 7804–7814, doi:10.18632/oncotarget.3158 (2015).

96 Stavreva, D. A. et al. Dynamics of chromatin accessibility and long-range interactions in response to glucocorticoid pulsing. Genome Res 25, 845–857, doi:10.1101/gr.184168.114 (2015).

97 Rickman, D. S. et al. Oncogene-mediated alterations in chromatin conformation. Proc Natl Acad Sci U S A 109, 9083–9088, doi:10.1073/pnas.1112570109 (2012).

98 Rubin, A. J. et al. Lineage-specific dynamic and pre-established enhancer-promoter contacts cooperate in terminal differentiation. Nat Genet 49, 1522–1528, doi:10.1038/ng.3935 (2017).

99 Stadhouders, R. et al. Transcription factors orchestrate dynamic interplay between genome topology and gene regulation during cell reprogramming. Nat Genet 50, 238–249, doi:10.1038/s41588-017-0030-7 (2018).

100 Muerdter, F. & Stark, A. Gene Regulation: Activation through Space. Curr Biol 26, R895–R898, doi:10.1016/j.cub.2016.08.031 (2016).

101 Zabidi, M. A. et al. Enhancer-core-promoter specificity separates developmental and housekeeping gene regulation. Nature 518, 556–559, doi:10.1038/nature13994 (2015).

102 Lambert, S. A. et al. The Human Transcription Factors. Cell 172, 650–665, doi:10.1016/j.cell.2018.01.029 (2018).

103 Beagrie, R. A. et al. Complex multi-enhancer contacts captured by genome architecture mapping. Nature 543, 519–524, doi:10.1038/nature21411 (2017).

104 Ing-Simmons, E. et al. Spatial enhancer clustering and regulation of enhancer-proximal genes by cohesin. Genome Res 25, 504–513, doi:10.1101/gr.184986.114 (2015).

105 Schmitt, A. D. et al. A Compendium of Chromatin Contact Maps Reveals Spatially Active Regions in the Human Genome. Cell Rep 17, 2042–2059, doi:10.1016/j.celrep.2016.10.061 (2016).

106 Dekker, J. & Misteli, T. Long-Range Chromatin Interactions. Cold Spring Harb Perspect Biol 7, a019356, doi:10.1101/cshperspect.a019356 (2015).

107 Krivega, I. & Dean, A. Chromatin looping as a target for altering erythroid gene expression. Ann N Y Acad Sci 1368, 31–39, doi:10.1111/nyas.13012 (2016).

108 Denker, A. & de Laat, W. The second decade of 3C technologies: detailed insights into nuclear organization. Genes Dev 30, 1357–1382, doi:10.1101/gad.281964.116 (2016).

109 Hnisz, D., Shrinivas, K., Young, R. A., Chakraborty, A. K. & Sharp, P. A. A Phase Separation Model for Transcriptional Control. Cell 169, 13–23, doi:10.1016/j.cell.2017.02.007 (2017).

110 Sabari, B. R. et al. Coactivator condensation at super-enhancers links phase separation and gene control. Science, doi:10.1126/science.aar3958 (2018).

