## Supplemental Figures for "Oncogenic Notch promotes long-range regulatory interactions within hyperconnected 3D cliques"

### Figure S1

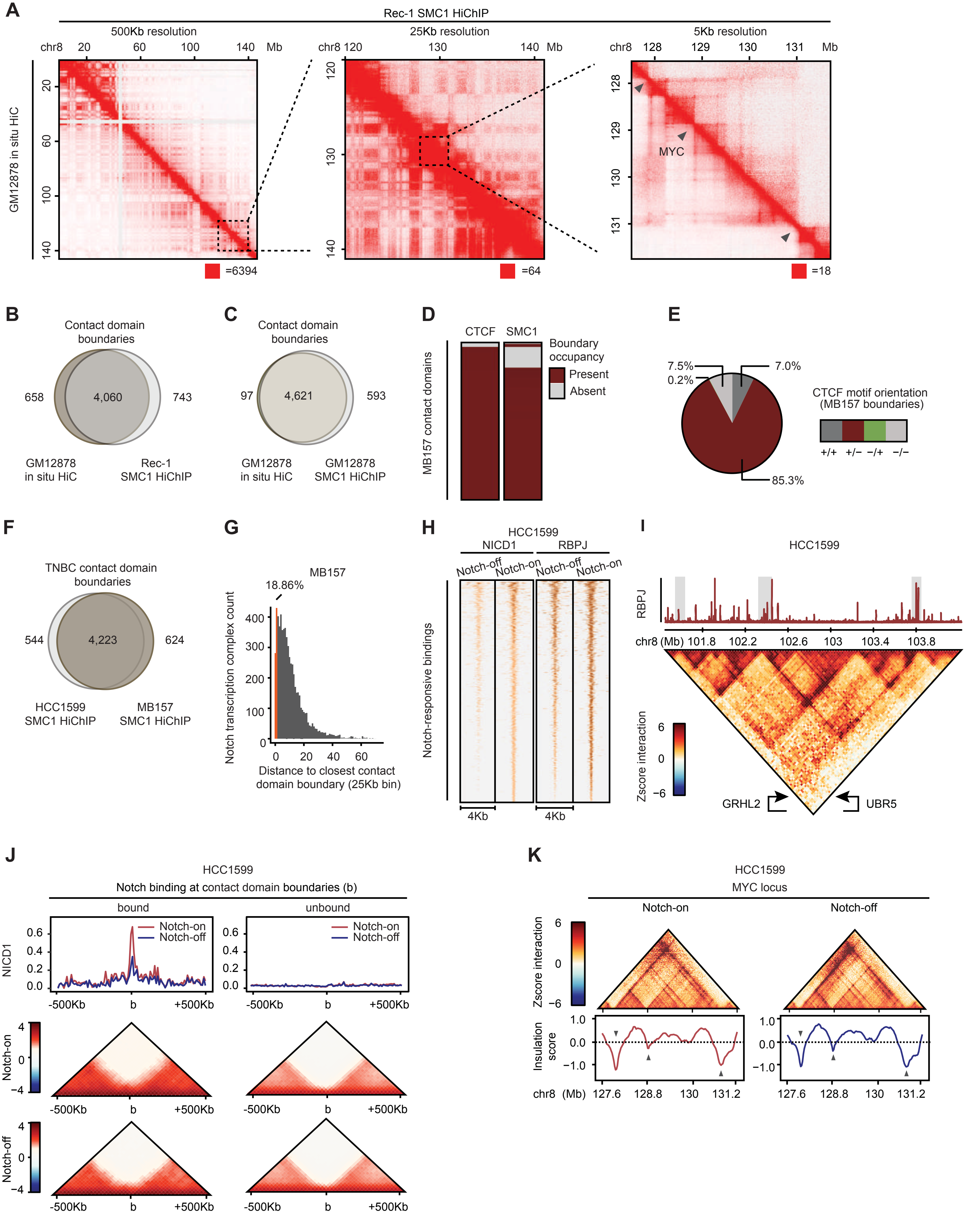
A
Rec-1 SMC1 HiChIP
500Kb resolution
25Kb resolution
5Kb resolution
GM12878 in situ HiC
MYC
=6394
=64
=18
B
Contact domain boundaries
658
4,060
743
GM12878 in situ HiC
Rec-1 SMC1 HiChIP
C
Contact domain boundaries
97
4,621
593
GM12878 in situ HiC
GM12878 SMC1 HiChIP
D
CTCF
SMC1
Boundary occupancy
Present
Absent
MB157 contact domains
E
7.5%
0.2%
7.0%
85.3%
CTCF motif orientation (MB157 boundaries)
+/+
+/−
−/+
−/−
F
TNBC contact domain boundaries
544
4,223
624
HCC1599 SMC1 HiChIP
MB157 SMC1 HiChIP
G
MB157
18.86%
Notch transcription complex count
Distance to closest contact domain boundary (25Kb bin)
H
HCC1599
NICD1
RBPJ
Notch-off
Notch-on
Notch-responsive bindings
4Kb
I
HCC1599
RBPJ
chr8 (Mb)
Zscore interaction
GRHL2
UBR5
J
HCC1599
Notch binding at contact domain boundaries (b)
bound
unbound
Notch-on
Notch-off
NICD1
-500Kb
b
+500Kb
K
HCC1599
MYC locus
Notch-on
Notch-off
Zscore interaction
Insulation score
chr8 (Mb)

**Figure S1. TNBC contact domain boundaries are mostly Notch-independent. Related to Figure 1.**

(A) Contact matrices depicting Rec-1 SMC1 HiChIP, upper half, and GM12878 in situ Hi-C, lower half, share contact domains at chromosome 8. Left: the whole chromosome, shown at 500 Kb resolution; middle: 120-140 Mb shown at 25 Kb resolution; right: 127.5 - 131.5 Mb *MYC* locus shown at 5 Kb resolution. The intensity of each pixel represents the normalized interaction frequency between two loci. Red square on the bottom right of each panel indicates maximum intensity. Gray arrows: boundaries demarcated by local minimum detection of insulation score.

(B,C) Venn diagrams comparing contact domain boundaries in (B) Rec-1 SMC1 HiChIP versus GM12878 *in situ* Hi-C (Fisher's exact p-value < 1E-15) (C) GM12878 *in situ* Hi-C versus SMC1 HiChIP (Fisher's exact p-value < 1E-04).

(D) Heatmap indicating the presence of CTCF and SMC1 on the boundaries of 2,317 MB157 contact domains.

(E) Pie chart showing the frequency of the four possible orientations of CTCF motif on contact domain boundary pairs depicts that the majority are in convergent (+/-) orientation.

(F) Venn diagram comparing contact domain boundaries in HCC1599 SMC1 HiChIP versus MB157 SMC1 HiChIP (Fisher's exact p-value < 1E-15).

(G) Histogram showing the distance between each Notch occupied site and its closest contact domain boundary in MB157. Orange mark: Notch binding within 3 x 25 Kb bins of a boundary.

(H) Heatmap of NICD1 and RBPJ occupancy shows 9,302 reproducible Notch binding events determined with IDR pipeline in HCC1599 with significant decrease (enrichR FDR < 0.05) in Notch-off (GSI) condition.

(I) Contact map (bottom) and genome-browser track of RBPJ ChIP-seq at *GRHL2* locus showing domain boundaries are enriched for Notch transcription complexes binding in HCC1599 similar to MB157.

(J) Metagene analyses (top) showing Notch occupancy, and pile-up plots (bottom) depicting aggregated Z-score interaction on HCC1599 domain boundaries in Notch-on (DMSO) and Notch-off (GSI) conditions where the overall differential boundary insulation scores are insignificant (Wilcoxon rank sum p-value > 0.11). Left: centered around 1,431 Notch-bound domain boundaries. Right: matching number of Notch-unbound boundaries.

(K) Contact map (top) and insulation profile (bottom) at *MYC* locus of HCC1599 in Notch-on (DMSO) and Notch-off (GSI) conditions showing intact domain boundaries. Gray arrows: boundaries demarcated by local minimum detection of insulation score.

#### Figure S2

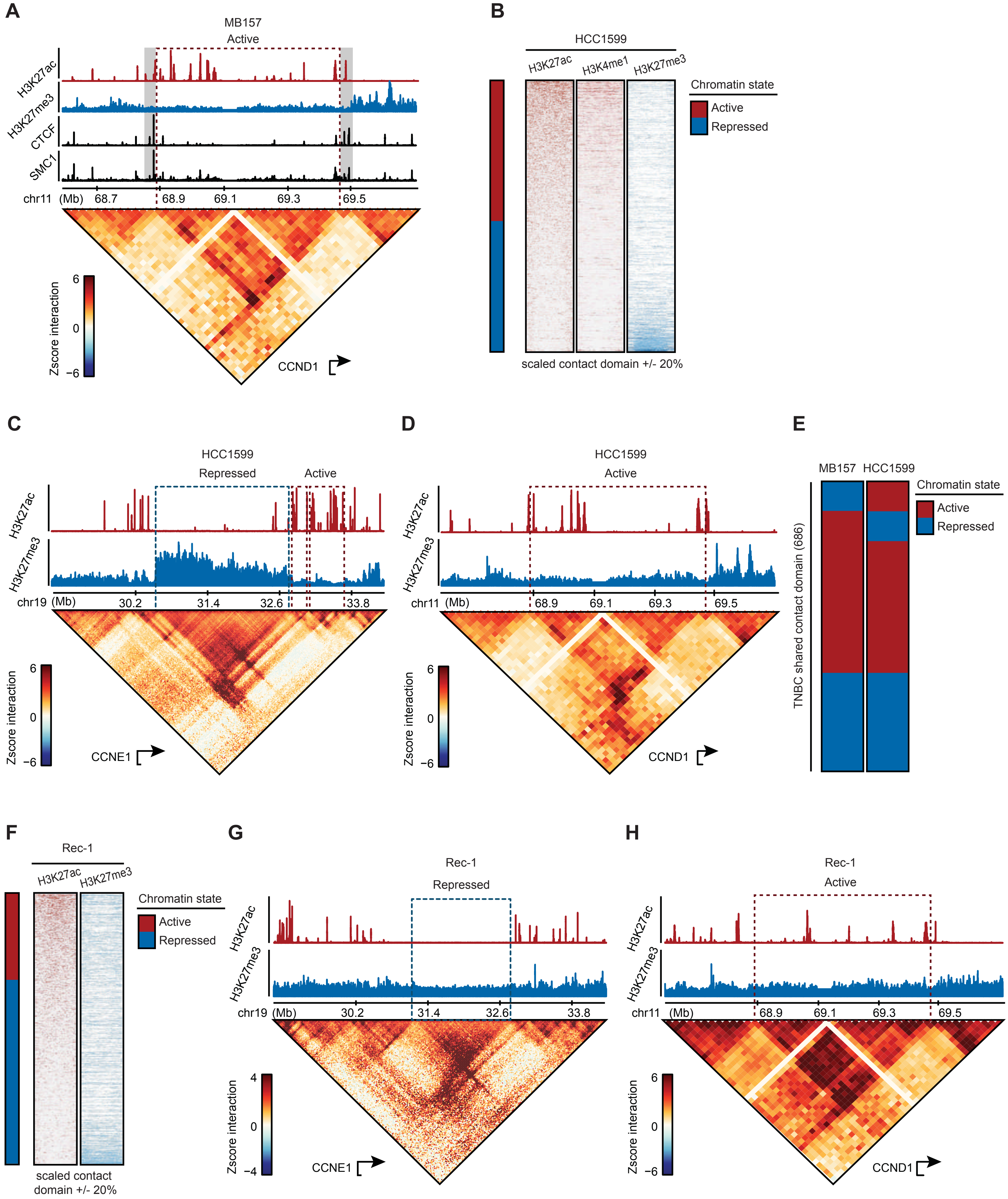
A
MB157
Active
H3K27ac
H3K27me3
CTCF
SMC1
chr11 (Mb) 68.7 68.9 69.1 69.3 69.5
Zscore interaction
6
0
−6
CCND1
B
HCC1599
H3K27ac
H3K4me1
H3K27me3
Chromatin state
Active
Repressed
scaled contact domain +/- 20%
C
HCC1599
Repressed
Active
H3K27ac
H3K27me3
chr19 (Mb) 30.2 31.4 32.6 33.8
Zscore interaction
6
0
−6
CCNE1
D
HCC1599
Active
H3K27ac
H3K27me3
chr11 (Mb) 68.9 69.1 69.3 69.5
Zscore interaction
6
0
−6
CCND1
E
MB157
HCC1599
Chromatin state
Active
Repressed
TNBC shared contact domain (686)
F
Rec-1
H3K27ac
H3K27me3
Chromatin state
Active
Repressed
scaled contact domain +/- 20%
G
Rec-1
Repressed
H3K27ac
H3K27me3
chr19 (Mb) 30.2 31.4 32.6 33.8
Zscore interaction
4
0
−4
CCNE1
H
Rec-1
Active
H3K27ac
H3K27me3
chr11 (Mb) 68.9 69.1 69.3 69.5
Zscore interaction
6
0
−6
CCND1

**Figure S2. Chromatin state aligns with contact domains in Notch-mutated cancers. Related to Figure 2.**

(A) Contact map (bottom) and genome browser tracks (top) in MB157 showing H3K27ac and H3K27me3 histone marks separated by contact domain boundaries with CTCF and SMC1 occupancy at *CCND1* locus. Red/blue dashed-line boxes: active/repressed domains. Gray boxes: marking CTCF and SMC1 binding events on the boundaries.

(B) Heatmap displaying normalized H3K27ac, H3K4me1 and H3K27me3 load within 1,709 contact domains with significant intradomain interactions in HCC1599. Each contact domain is categorized into active or repressed based on the differential H3K27ac and H3K27me3 total level and sorted in descending order.

(C) Contact map (bottom) and genome browser tracks of H3K27ac and H3K27me3 (top) in HCC1599 showing contact domain boundaries and chromatin state shared between MB157 and HCC1599 at *CCNE1* locus. Red/blue dashed-line boxes: active/repressed domains.

(D) Contact map (bottom) and genome browser tracks (top) in HCC1599 showing H3K27ac and H3K27me3 histone marks separated by contact domain boundaries at *CCND1* locus.

(E) Heatmap showing chromatin state of 686 contact domains shared between MB157 and HCC1599 cells (Fisher's exact p-value < 1E-15).

(F) Same heatmap as in (B) showing 1,523 contact domains in Rec-1.

(G) *CCNE1* locus as (C) in Rec-1

(H) *CCND1* locus as (D) in Rec-1.

#### Figure S3

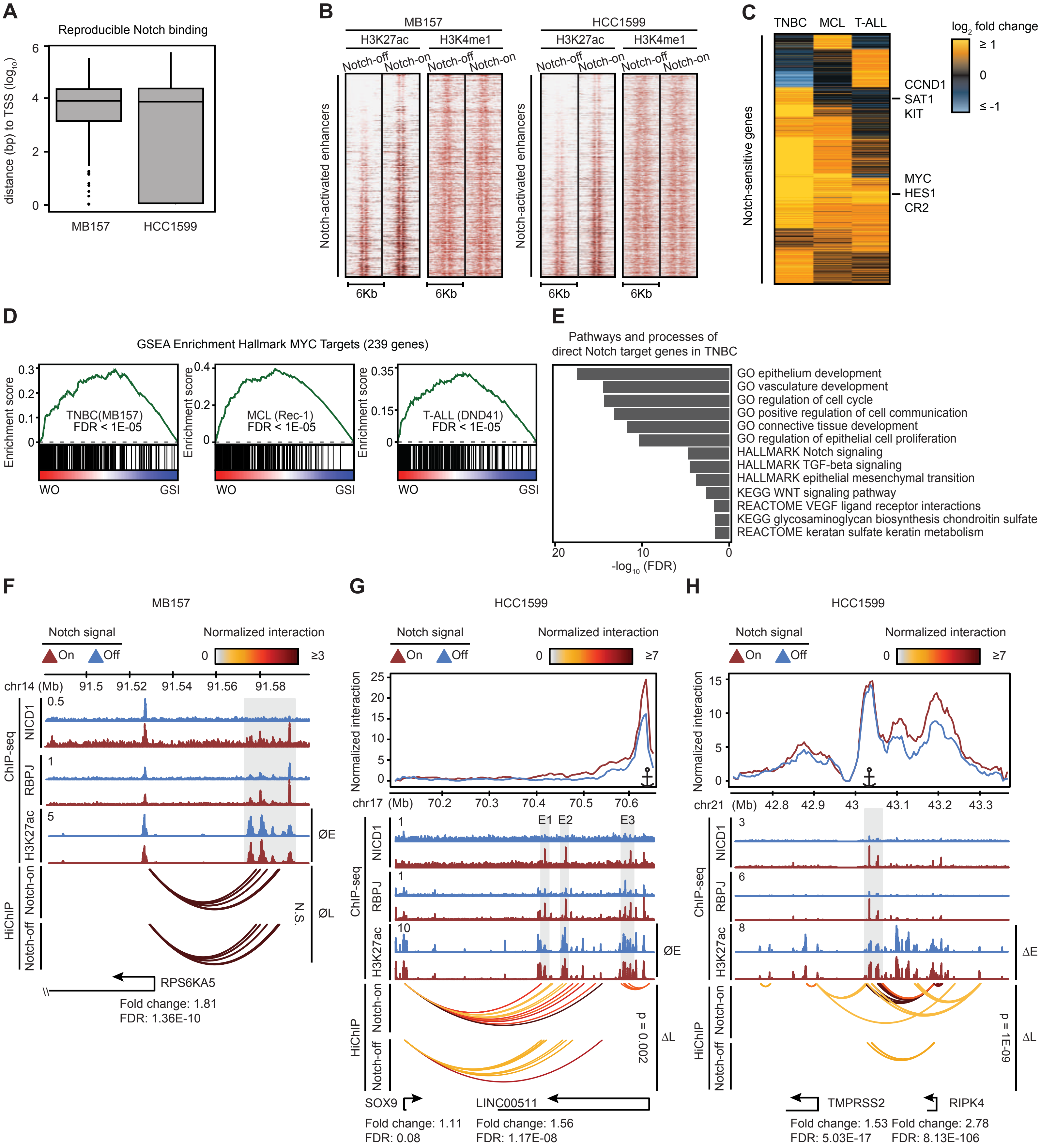
A
Reproducible Notch binding
distance (bp) to TSS (log10)
MB157
HCC1599
B
MB157
HCC1599
H3K27ac
H3K4me1
Notch-off
Notch-on
Notch-activated enhancers
6Kb
C
TNBC
MCL
T-ALL
log2 fold change
≥ 1
0
≤ -1
Notch-sensitive genes
CCND1
SAT1
KIT
MYC
HES1
CR2
D
GSEA Enrichment Hallmark MYC Targets (239 genes)
Enrichment score
TNBC(MB157)
FDR < 1E-05
MCL (Rec-1)
FDR < 1E-05
T-ALL (DND41)
FDR < 1E-05
WO
GSI
E
Pathways and processes of
direct Notch target genes in TNBC
GO epithelium development
GO vasculature development
GO regulation of cell cycle
GO positive regulation of cell communication
GO connective tissue development
GO regulation of epithelial cell proliferation
HALLMARK Notch signaling
HALLMARK TGF-beta signaling
HALLMARK epithelial mesenchymal transition
KEGG WNT signaling pathway
REACTOME VEGF ligand receptor interactions
KEGG glycosaminoglycan biosynthesis chondroitin sulfate
REACTOME keratan sulfate keratin metabolism
-log10 (FDR)
F
MB157
Notch signal
On
Off
Normalized interaction
0
≥3
chr14 (Mb)
91.5
91.52
91.54
91.56
91.58
ChIP-seq
NICD1
RBPJ
H3K27ac
HiChIP
Notch-on
Notch-off
ØE
ØL
N.S.
RPS6KA5
Fold change: 1.81
FDR: 1.36E-10
G
HCC1599
≥7
Normalized interaction
chr17 (Mb)
70.2
70.3
70.4
70.5
70.6
E1
E2
E3
ØE
ΔL
p = 0.002
SOX9
LINC00511
Fold change: 1.11
FDR: 0.08
Fold change: 1.56
FDR: 1.17E-08
H
HCC1599
chr21 (Mb)
42.8
42.9
43
43.1
43.2
43.3
ΔE
ΔL
p = 1E-09
TMPRSS2
RIPK4
Fold change: 1.53
FDR: 5.03E-17
Fold change: 2.78
FDR: 8.13E-106

**Figure S3. Notch activates TNBC distal enhancers. Related to Figure 3**

(A) Boxplot showing the linear genomic distance between Notch binding sites and their closest Ensembl annotated transcription start site (TSS) in $\log_{10}$ scale.

(B) Heatmap showing H3K27ac and H3K4me1 level at Notch-activated enhancers centered around Notch binding sites +/- 3 kb flanking regions in MB157 (left) and HCC1599 (right). Each row is an enhancer sorted by $\log_2$ fold change of H3K27ac level in Notch-on (GSI-washout) versus Notch-off (GSI).

(C) Heatmap displaying common and unique Notch-sensitive genes in TNBC (HCC1599 and MB157), MCL (Rec-1) and T-ALL (DND41). Each row is a gene grouped by K-means clustering on differential expression - $\log_2$ fold change of Notch-on (GSI-washout) vs Notch-off (GSI).

(D) GSEA analysis showing the enrichment of genes in the MSigDB Hallmark MYC targets in Notch-on (GSI-washout) vs Notch-off (GSI) in MB157, Rec-1 and DND41 (permutation FDR < 1E-05).

(E) Selected GO terms and pathways enriched with direct Notch target genes in TNBC. MSigDB was used for analysis of functional gene annotation.

(F) Notch as a final transcriptional trigger (example of mode d). Notch-bound but Notch-insensitive loops (ØL) linking Notch-bound but Notch-insensitive enhancers (ØE) to *PRS6KA5* in MB157. ChIP-seq tracks showing Notch-sensitive NICD1 and RBPJ occupancy and Notch-insensitive H3K27ac level marked with gray box. HiChIP arcs displaying Notch-insensitive normalized significant interactions between enhancers and promoters in Notch-on (DMSO, top) and Notch-off (GSI, bottom) in MB157 (paired t-test p-value > 0.05). Bottom track indicating Ensembl gene position of *RPS6KA5* and its expression fold change and FDR determined by DESeq2.

(G) Same locus as in Figure 3I in HCC1599.

(H) Same locus as Figure 3J in HCC1599.

### Figure S4

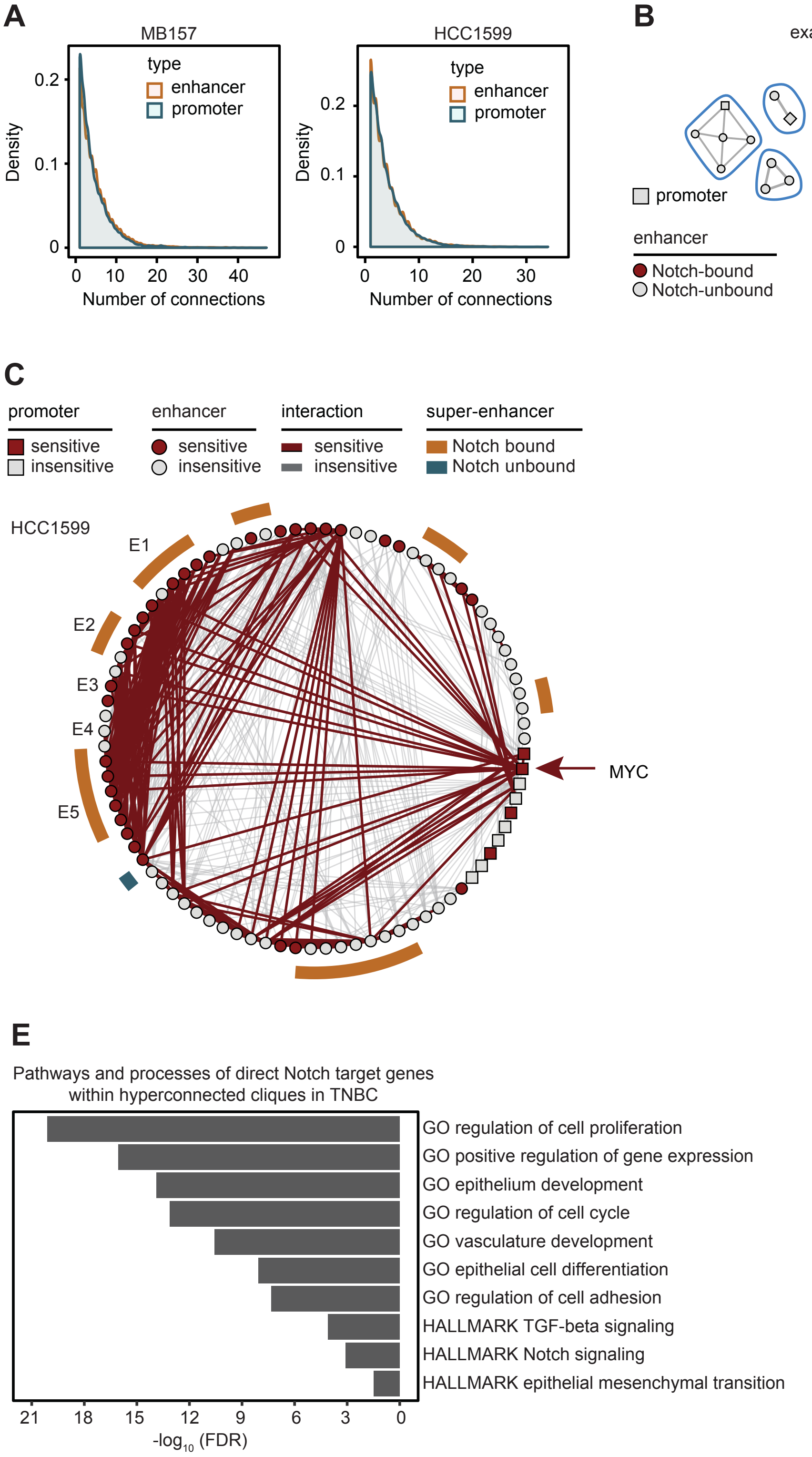
A
MB157
HCC1599
type
enhancer
promoter
Density
Number of connections
B
example of 4E (MB157)
example of 4E (HCC1599)
promoter
enhancer
Notch-bound
Notch-unbound
C
promoter
sensitive
insensitive
enhancer
sensitive
insensitive
interaction
sensitive
insensitive
super-enhancer
Notch bound
Notch unbound
HCC1599
E1
E2
E3
E4
E5
MYC
D
MB157
HCC1599
p < 1E-15
Number of enhancer per promoter
ΔE + ΔL
øE + ΔL
ΔE + øL
øE + øL
Notch dependency
E
Pathways and processes of direct Notch target genes within hyperconnected cliques in TNBC
GO regulation of cell proliferation
GO positive regulation of gene expression
GO epithelium development
GO regulation of cell cycle
GO vasculature development
GO epithelial cell differentiation
GO regulation of cell adhesion
HALLMARK TGF-beta signaling
HALLMARK Notch signaling
HALLMARK epithelial mesenchymal transition
-log10 (FDR)

**Figure S4. Notch target genes within hyperconnected cliques associate with key pathways in TNBC. Related to Figure 4.**

(A) Density plots showing the number of connections to each enhancer or promoter in MB157 (left) and HCC1599 (right).

(B) Three randomly selected cliques with or without Notch-bound enhancers in MB157 (left) and HCC1599 (right) from Figure 4E emphasizing on differences in their connectivity.

(C) Circos plot showing the clique associated with *MYC* in HCC1599. Red-marked circle (square) and line depicting Notch-sensitive enhancer (promoter) and significant long-range interactions, respectively. E1 to E5 mark groups of enhancers in descending linear genomic distance to *MYC* promoter located within the *MYC* 5' contact domain.

(D) Barplots depicting the average ± SEM number of enhancers connecting each Notch-sensitive gene with combination of Notch-bound and -promoted loops (ΔL), Notch-bound and -activated enhancers (ΔE), Notch-bound but Notch-insensitive loops (ØL), and Notch-bound but Notch-insensitive enhancers (ØE) in MB157 (left) and HCC1599 (right) (Wilcoxon rank sum p-value < 1E-15).

(E) Selected GO terms and pathways enriched with direct Notch target genes within hyperconnected cliques in TNBC. MSigDB was used for analysis of functional gene annotation.

### Figure S5

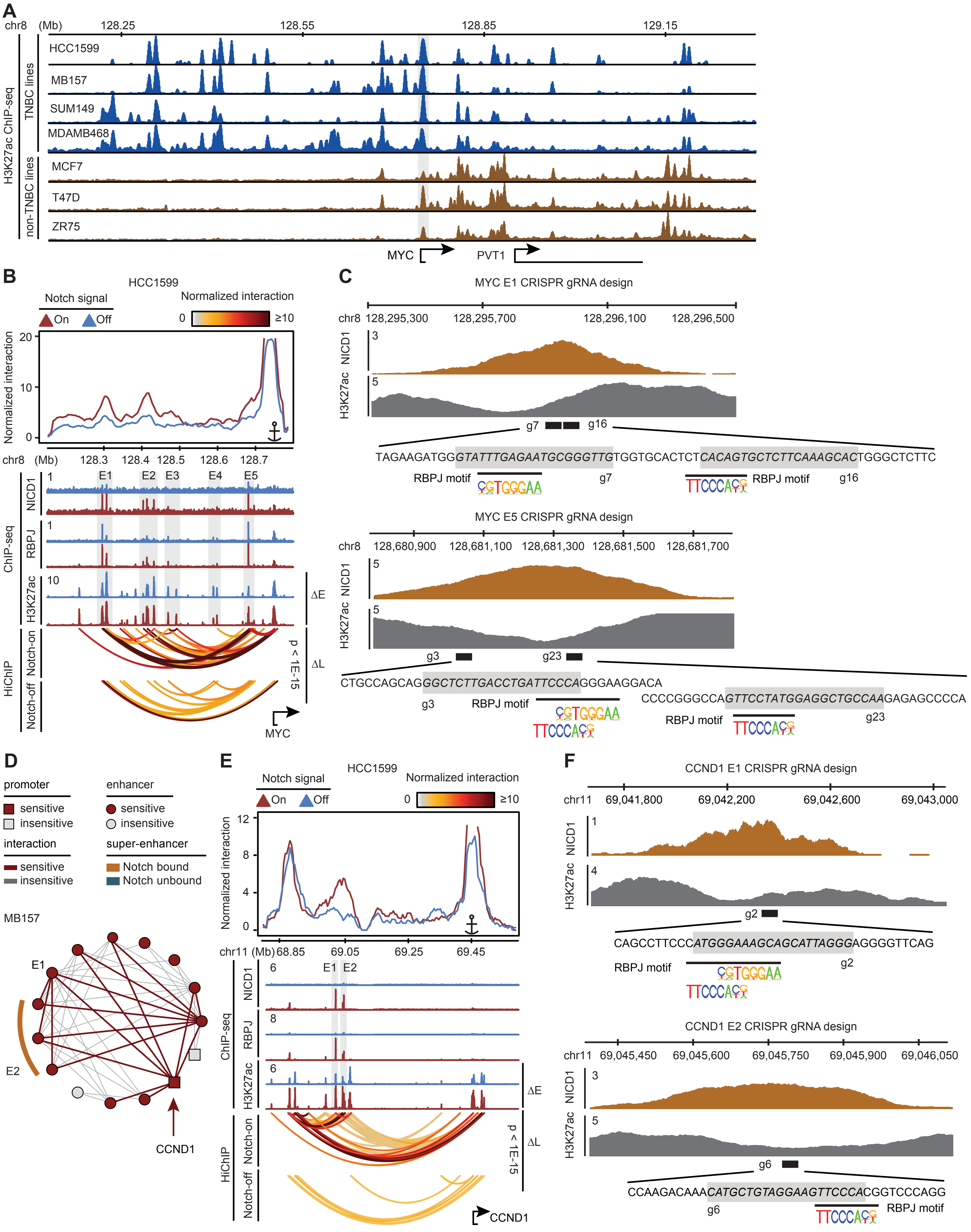
A
chr8 (Mb) 128.25 128.55 128.85 129.15
H3K27ac ChIP-seq
TNBC lines
HCC1599
MB157
SUM149
MDAMB468
non-TNBC lines
MCF7
T47D
ZR75
MYC
PVT1
B
HCC1599
Notch signal
On Off
Normalized interaction
0 ≥10
Normalized interaction
20
10
0
chr8 (Mb) 128.3 128.4 128.5 128.6 128.7
ChIP-seq
NICD1
RBPJ
H3K27ac
E1 E2 E3 E4 E5
1
1
10
ΔE
HiChIP
Notch-on
Notch-off
p < 1E-15
ΔL
MYC
C
MYC E1 CRISPR gRNA design
chr8 128,295,300 128,295,700 128,296,100 128,296,500
NICD1
H3K27ac
3
5
g7 g16
TAGAAGATGGGTATTTGAGAATGCGGGTTGTGGTGCACTCTCACAGTGCTCTTCAAAGCACTGGGCTCTTC
RBPJ motif
g7
RBPJ motif
g16
MYC E5 CRISPR gRNA design
chr8 128,680,900 128,681,100 128,681,300 128,681,500 128,681,700
NICD1
H3K27ac
5
5
g3 g23
CTGCCAGCAGGGCTCTTGACCTGATTCCCAGGGAAGGACA
g3
RBPJ motif
CCCCGGGCCAGTTCCTATGGAGGCTGCCAAGAGAGCCCCA
RBPJ motif
g23
D
promoter
sensitive
insensitive
enhancer
sensitive
insensitive
interaction
sensitive
insensitive
super-enhancer
Notch bound
Notch unbound
MB157
E1
E2
CCND1
E
HCC1599
Notch signal
On Off
Normalized interaction
0 ≥10
Normalized interaction
12
8
4
0
chr11 (Mb) 68.85 69.05 69.25 69.45
ChIP-seq
NICD1
RBPJ
H3K27ac
6
8
6
E1 E2
ΔE
HiChIP
Notch-on
Notch-off
p < 1E-15
ΔL
CCND1
F
CCND1 E1 CRISPR gRNA design
chr11 69,041,800 69,042,200 69,042,600 69,043,000
NICD1
H3K27ac
1
4
g2
CAGCCTTCCCATGGGAAAGCAGCATTAGGGAGGGGTTCAG
RBPJ motif
g2
CCND1 E2 CRISPR gRNA design
chr11 69,045,450 69,045,600 69,045,750 69,045,900 69,046,050
NICD1
H3K27ac
3
5
g6
CCAAGACAAACATGCTGTAGGAAGTTCCCACGGTCCCAGG
g6
RBPJ motif

**Figure S5: Targeting *MYC* and *CCND1* Notch-dependent enhancers in TNBC. Related to Figure 5.**

(A) Genome-browser tracks showing difference between the H3K27ac level of TNBC and non-TNBC breast cancer lines at *MYC* locus.

(B) Same locus as in Figure 5A in HCC1599.

(C) NICD1 binding and H3K27ac level at *MYC* E1 and E5 enhancers in MB157 cells. Consensus RBPJ logos are aligned with CRISPR-targeted RBPJ motifs determined by FIMO (p-value < 1E-03) in each enhancer. The sgRNAs sequences are italicized and their positions are boxed.

(D) The clique associated with *CCND1* in MB157.

(E) Same locus as in Figure 5F in HCC1599.

(F) NICD1 binding and H3K27ac level at *CCND1* E1 and E2 enhancers in MB157 cells. Consensus RBPJ logos are aligned with CRISPR-targeted RBPJ motifs determined by FIMO (p-value < 1E-03) in each enhancer. The sgRNAs sequences are italicized and their positions are boxed.

### Figure S6

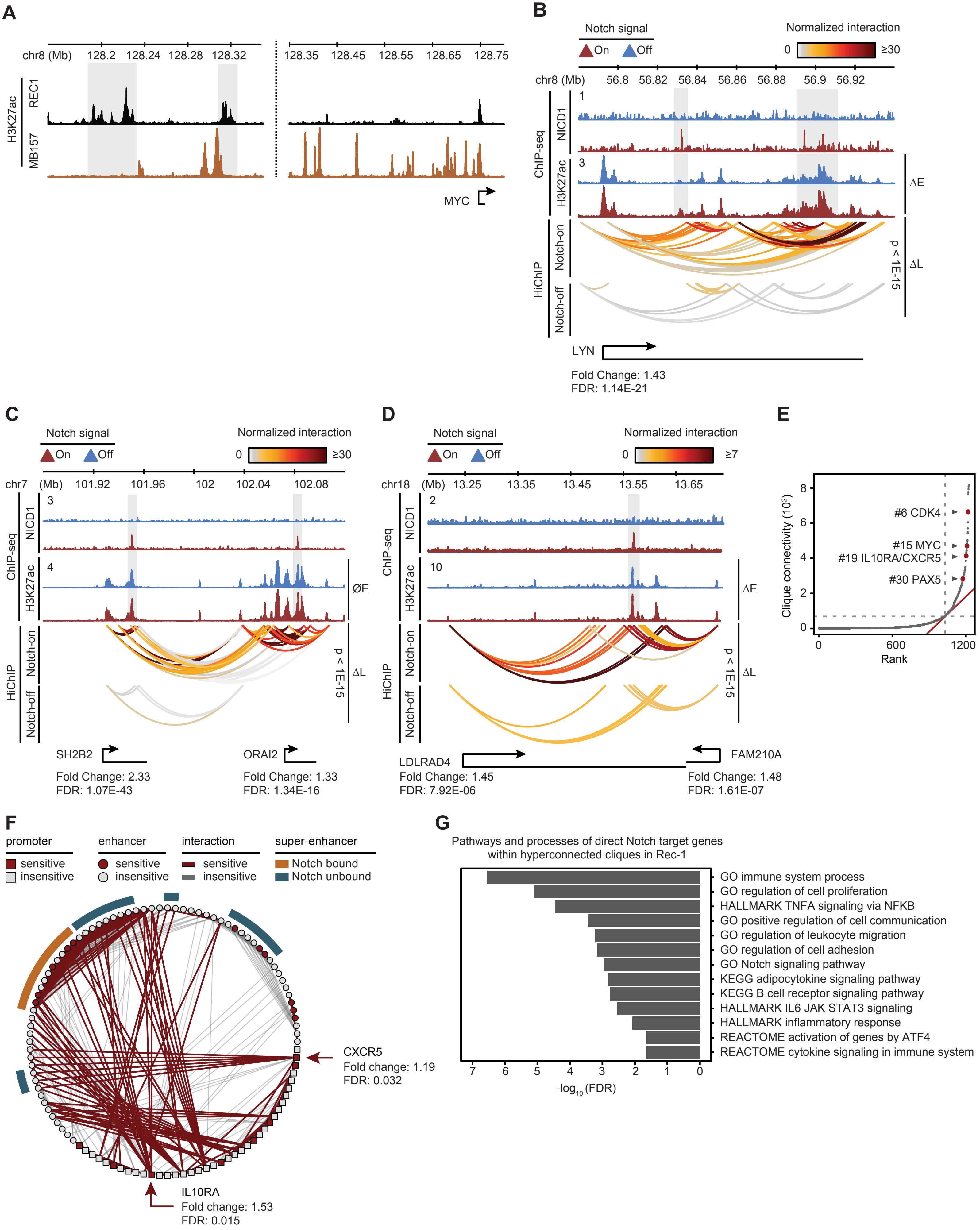

A
chr8 (Mb)
128.2
128.24
128.28
128.32
128.35
128.45
128.55
128.65
128.75
H3K27ac
REC1
MB157
MYC
B
Notch signal
On
Off
Normalized interaction
0
≥30
chr8 (Mb)
56.8
56.82
56.84
56.86
56.88
56.9
56.92
ChIP-seq
NICD1
H3K27ac
HiChIP
Notch-on
Notch-off
ΔE
ΔL
p < 1E-15
LYN
Fold Change: 1.43
FDR: 1.14E-21
C
chr7 (Mb)
101.92
101.96
102
102.04
102.08
ØE
SH2B2
Fold Change: 2.33
FDR: 1.07E-43
ORAI2
Fold Change: 1.33
FDR: 1.34E-16
D
≥7
chr18 (Mb)
13.25
13.35
13.45
13.55
13.65
LDLRAD4
Fold Change: 1.45
FDR: 7.92E-06
FAM210A
Fold Change: 1.48
FDR: 1.61E-07
E
Clique connectivity (10²)
Rank
#6 CDK4
#15 MYC
#19 IL10RA/CXCR5
#30 PAX5
F
promoter
sensitive
insensitive
enhancer
interaction
super-enhancer
Notch bound
Notch unbound
CXCR5
Fold change: 1.19
FDR: 0.032
IL10RA
Fold change: 1.53
FDR: 0.015
G
Pathways and processes of direct Notch target genes within hyperconnected cliques in Rec-1
GO immune system process
GO regulation of cell proliferation
HALLMARK TNFA signaling via NFKB
GO positive regulation of cell communication
GO regulation of leukocyte migration
GO regulation of cell adhesion
GO Notch signaling pathway
KEGG adipocytokine signaling pathway
KEGG B cell receptor signaling pathway
HALLMARK IL6 JAK STAT3 signaling
HALLMARK inflammatory response
REACTOME activation of genes by ATF4
REACTOME cytokine signaling in immune system
-log₁₀ (FDR)

**Figure S6. Examples of Notch-promoted loops in MCL. Related to Figure 6.**

(A) H3K27ac ChIP-seq tracks showing MCL-specific (Rec-1) and TNBC-specific (MB157) *MYC* enhancers. Gray box: MCL-specific enhancers.

(B) Notch-promoted loops (ΔL) linking Notch-activated enhancers (ΔE) to *LYN* in Rec-1. ChIP-seq tracks showing Notch-sensitive NICD1 occupancy, and Notch-sensitive H3K27ac level. HiChIP arcs displaying normalized significant interactions of *LYN* promoter to distal enhancers, and enhancer-enhancer interactions in Notch-on (DMSO, top) and Notch-off (GSI, bottom) (paired t-test p-value $< 1E-15$). Bottom track indicating *LYN* Ensembl gene position and its expression fold change and FDR as determined by DESeq2.

(C) Notch-promoted loops (ΔL) linking Notch-bound but Notch-insensitive enhancers (ØE) to *SH2B2* promoter in Rec-1. Tracks' descriptions as in panel (B).

(D) Notch-promoted loops (ΔL) enabling spatial co-regulation of *FAM210A* and *LDLRAD4* genes in MCL through shared Notch-activated enhancers (ΔE). Tracks' descriptions as in panel (B).

(E) Distribution of 3D cliques connectivity in Rec-1 plotted in an ascending order. Example cliques are marked and named with their representative Notch-sensitive genes.

(F) Circos plot showing the clique associated with *IL10RA*, one of the known Notch target genes in Rec-1. Red-marked circle (square) and line depicting Notch-sensitive enhancer (promoter) and significant long-range interactions, respectively.

(G) Selected GO terms and pathways enriched with direct Notch target genes within hyperconnected cliques in Rec-1. MSigDB was used for analysis of functional gene annotation.

### Figure S7

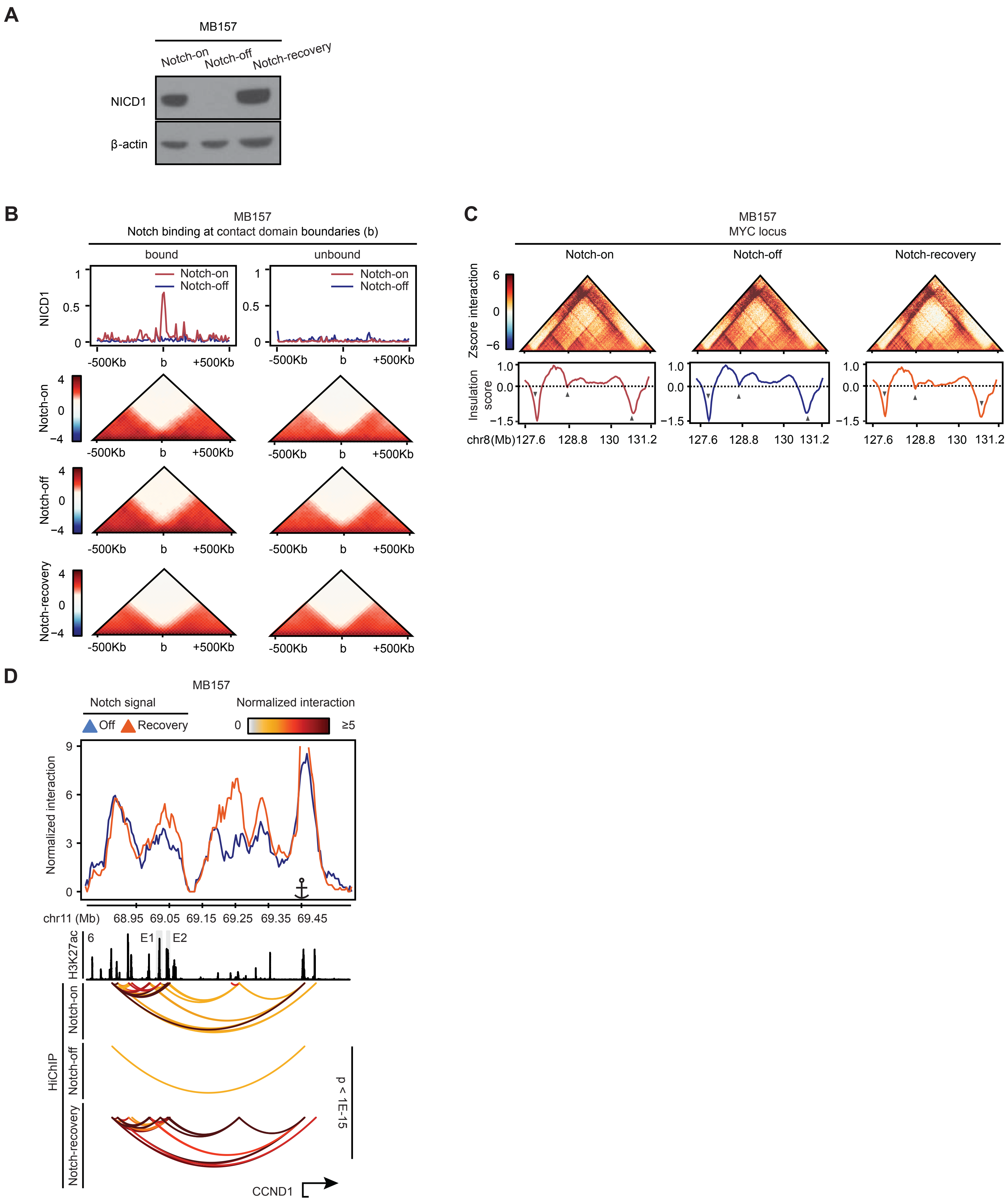
A
MB157
Notch-on
Notch-off
Notch-recovery
NICD1
β-actin
B
MB157
Notch binding at contact domain boundaries (b)
bound
unbound
NICD1
Notch-on
Notch-off
Notch-recovery
-500Kb
b
+500Kb
C
MB157
MYC locus
Notch-on
Notch-off
Notch-recovery
Zscore interaction
Insulation score
chr8(Mb)
127.6
128.8
130
131.2
D
MB157
Notch signal
Off
Recovery
Normalized interaction
0
≥5
Normalized interaction
chr11 (Mb)
68.95
69.05
69.15
69.25
69.35
69.45
H3K27ac
E1
E2
HiChIP
Notch-on
Notch-off
Notch-recovery
p < 1E-15
CCND1

**Figure S7. Contact domains are invariant in Notch recovery. Related to Figure 7.**

(A) Western blot validating recovery of NICD1 in GSI-washout. $\beta$-actin is loading control.

(B) Metagene analyses (top) showing Notch occupancy, and pile-up plots (bottom) in MB157 contact domain boundaries in Notch-on (DMSO), Notch-off (GSI), and Notch-recovery (GSI-washout) (Wilcoxon rank sum p-value > 0.15). Left: centered around 1,003 Notch-bound domain boundaries. Right: matching number of Notch-unbound boundaries.

(C) Contact map (top) and insulation profile (bottom) at *MYC* locus showing contact domain boundaries are similar in Notch-on (DMSO), Notch-off (GSI) and Notch-recovery (GSI-washout) conditions. Gray arrows: boundaries identified by local minimum detection of insulation score.

(D) Notch recovery rescues interactions at *CCND1* locus in MB157. Top panel: virtual 4C plot depicting the normalized interaction frequency from *CCND1* promoter viewpoint. ChIP-seq tracks showing H3K27ac level. HiChIP arcs displaying normalized significant interactions between *CCND1* promoter and distal enhancers in Notch-on (top, DMSO), Notch-off (middle, GSI), and Notch-recovery (bottom, GSI-washout).
